# An integrated mechanistic and data-driven computational model predicts cell responses to high- and low-affinity EGFR ligands

**DOI:** 10.1101/2023.06.25.543329

**Authors:** Paul J. Myers, Sung Hyun Lee, Matthew J. Lazzara

## Abstract

The biophysical properties of ligand binding heavily influence the ability of receptors to specify cell fates. Understanding the rules by which ligand binding kinetics impact cell phenotype is challenging, however, because of the coupled information transfers that occur from receptors to downstream signaling effectors and from effectors to phenotypes. Here, we address that issue by developing an integrated mechanistic and data-driven computational modeling platform to predict cell responses to different ligands for the epidermal growth factor receptor (EGFR). Experimental data for model training and validation were generated using MCF7 human breast cancer cells treated with the high- and low-affinity ligands epidermal growth factor (EGF) and epiregulin (EREG), respectively. The integrated model captures the unintuitive, concentration-dependent abilities of EGF and EREG to drive signals and phenotypes differently, even at similar levels of receptor occupancy. For example, the model correctly predicts the dominance of EREG over EGF in driving a cell differentiation phenotype through AKT signaling at intermediate and saturating ligand concentrations and the ability of EGF and EREG to drive a broadly concentration-sensitive migration phenotype through cooperative ERK and AKT signaling. Parameter sensitivity analysis identifies EGFR endocytosis, which is differentially regulated by EGF and EREG, as one of the most important determinants of the alternative phenotypes driven by different ligands. The integrated model provides a new platform to predict how phenotypes are controlled by the earliest biophysical rate processes in signal transduction and may eventually be leveraged to understand receptor signaling system performance depends on cell context.

**One-sentence summary:** Integrated kinetic and data-driven EGFR signaling model identifies the specific signaling mechanisms that dictate cell responses to EGFR activation by different ligands.

## INTRODUCTION

The epidermal growth factor receptor (EGFR) initiates cell signaling processes that are required for normal tissue development but can be coopted by cancers to promote tumor progression (*1–4*). The cellular consequences of EGFR activation are dictated in part by the biophysical properties of the specific ligand that binds the receptor, but the rules that map ligand properties to cell fates are incompletely understood. Determining those rules is challenging because they span two tiers of the signal transduction process: signaling network activation by the receptor and cellular interpretation of multivariate effector signals to make phenotypic decisions.

There is some mechanistic understanding of the basis by which different EGFR ligands regulate cell outcomes. The low-affinity EGFR ligands epigen (EPGN) and epiregulin (EREG) promote weak but long-lived EGFR dimers, whereas the high-affinity ligand EGF promotes stronger, short-lived dimers (*5*). Additionally, at comparable levels of receptor saturation, EPGN- or EREG-bound EGFR undergoes slower endocytosis and degradation than EGF-bound EGFR (*6, 7*), and EREG and EPGN induce greater GRB2-EGFR association than EGF. (*8*). Collectively, these effects result in EREG and EPGN promoting more protracted ERK signaling. This is a proposed explanation for the ability of low-affinity ligands to induce cell differentiation more potently than EGF (*5, 7, 9, 10*), though the effects of other pathways on phenotypes have not been fully explored. The strength and duration of EGFR-initiated signaling is also well-known to be regulated by phosphatase expression and localization (*2, 11–13*) and feedbacks mediated by ERK, PI3K, RAS GTPase-activating proteins (RASGAP) (*2, 14–16*). Each of these processes may be influenced by the biophysical properties of the specific ligand that binds EGFR.

Computational systems biology models provide a powerful framework for quantitatively predicting how perturbations at different levels of a signal transduction cascade propagate downstream. Mechanistic, differential equation-based models can explain the dynamics of EGFR and effector pathway activation in terms of parameters describing receptor homo-and hetero-dimerization (*17–22*), receptor endocytosis and downstream trafficking (*23–25*), regulation by protein tyrosine phosphatases (PTPs) (*11, 13*), signaling amplification and feedback loops (*26–32*), and the response of the EGFR signaling network to mutations and pharmacological interventions (*33–36*). Mechanistic models are generally inappropriate, however, for predicting phenotypes, which are not governed by species conservation laws or physics-based constitutive equations (*37*). Rather, the quantitative relationships that govern cell phenotypes are best identified by data-driven models based on high-dimensional feature data at the transcriptomic or (phospho)proteomic level (*37*). Such classification or regression models can distill the network to identify key independent variables that dictate cell outcomes in response to EGFR signaling (*38–41*). A natural extension of prior modeling approaches would be a unified framework that simulates signaling dynamics and resultant phenotypes in response to EGFR activation. Some prior work touches on these concepts. For example, a large-scale, ordinary differential equation (ODE) model of ErbB signaling predicts cell viability responses to pharmacological perturbations using Cancer Cell Line Encyclopedia (CCLE) data from over 100 cancer cell lines(*36*); however, model training relied solely on cell death phenotype measurements. A model of β-cell insulin secretion—trained on dynamic mass spectrometry-based metabolite and insulin measurements—integrates ODEs with a partial least squares regression (PLSR) data-driven model; using ODE-predicted metabolite levels as PLSR inputs, this integrated model predicts β-cell insulin secretion in response to glucose treatment (*42*). Notably, however, this model focuses exclusively on central carbon metabolism rather than RTK-regulated phosphoprotein signaling dynamics and does not address the fundamental question of phenotype determination by different agonists for the same receptor.

Here, we develop an integrated model of EGFR signaling to predict how changes to EGFR ligand-receptor binding properties alter signaling network activation to impact cell outcomes. The model comprises a set of mechanistic ordinary differential equations that predict EGFR-ERK-AKT network dynamic response to EGFR ligation and a coupled PLSR model that predicts phenotypes from mechanistic model outputs. The model is trained on signaling and phenotype measurements in MCF7 human breast cancer cells, a useful background for studying the effects of high- and low-affinity EGFR ligands (*5*). The trained integrated model captures the concentration-dependent abilities of EGF and EREG to drive different phenotypes at similar levels of receptor occupancy in ways that involve effector pathway cooperation or dominance of one pathway relative to another. Ultimately, this new computational framework enables the calculation of model sensitivities from biophysical properties all the way to phenotypic outcomes, and the approach is readily extendable to other signaling systems and other cell contexts.

## RESULTS

### Differential signaling and cellular responses induced by EGF and EREG

To construct an integrated model of cell responses to EGFR activation, we combined a coupled ODE model describing EGFR-ERK-AKT signaling dynamics with a data-driven PLSR model that predicts cell phenotypes based on dynamic signaling protein inputs (**Fig. 1**). We trained the model using data gathered from MCF7 cells, which exhibit differential signaling and phenotypes when treated with high- or low-affinity EGFR ligands (*5, 43*). MCF7 cells were treated with three concentrations of EGF or EREG, and signaling dynamics, differentiation, and migration were measured. Ligands were used at concentrations near receptor saturation, half-saturation, and sub-saturation based on their affinities (∼1.6 nM for EGF, ∼76 nM for EREG; see *Materials and Methods*). Signaling data were gathered for the first 90 min of response to ligands, based on a desire for the ODE model to capture only early signaling responses, which can be predictive of phenotypic responses at much longer times (*5, 43, 44*). Consistent with prior studies (*5*), saturating EGF induced more transient phosphorylation of AKT and ERK than saturating EREG (**Fig. 2A**). EGFR phosphorylation was similar for the two ligands over the times assessed. At half-saturation, EGF and EREG produced similar ERK and AKT phosphorylation dynamics, with EREG producing only slightly higher AKT phosphorylation. At the sub-saturating doses, EGF induced notably more sustained ERK phosphorylation than EREG. Sustained AKT and ERK activation by larger EREG concentrations coincided with strong MCF7 differentiation, indicated by neutral lipid accumulation (Oil Red O staining), which was much less apparent with EGF or low-dose EREG (**Fig. 2B**). Consistent with prior observations (*43*), however, sub-saturating EGF induced strong migratory responses; the saturating dose of EREG similarly produced a strong migratory response (**Fig. 2C**). These responses again mostly coincided with more sustained activation of both ERK and AKT, and the overall trend of EREG as a driver of more sustained ERK and AKT activation was also apparent 24 hr after ligand treatment (**Fig. 2D**). It is unclear from the signaling data alone, however, if one pathway controls phenotypes more than the other or if they cooperate.

**Figure 1.**
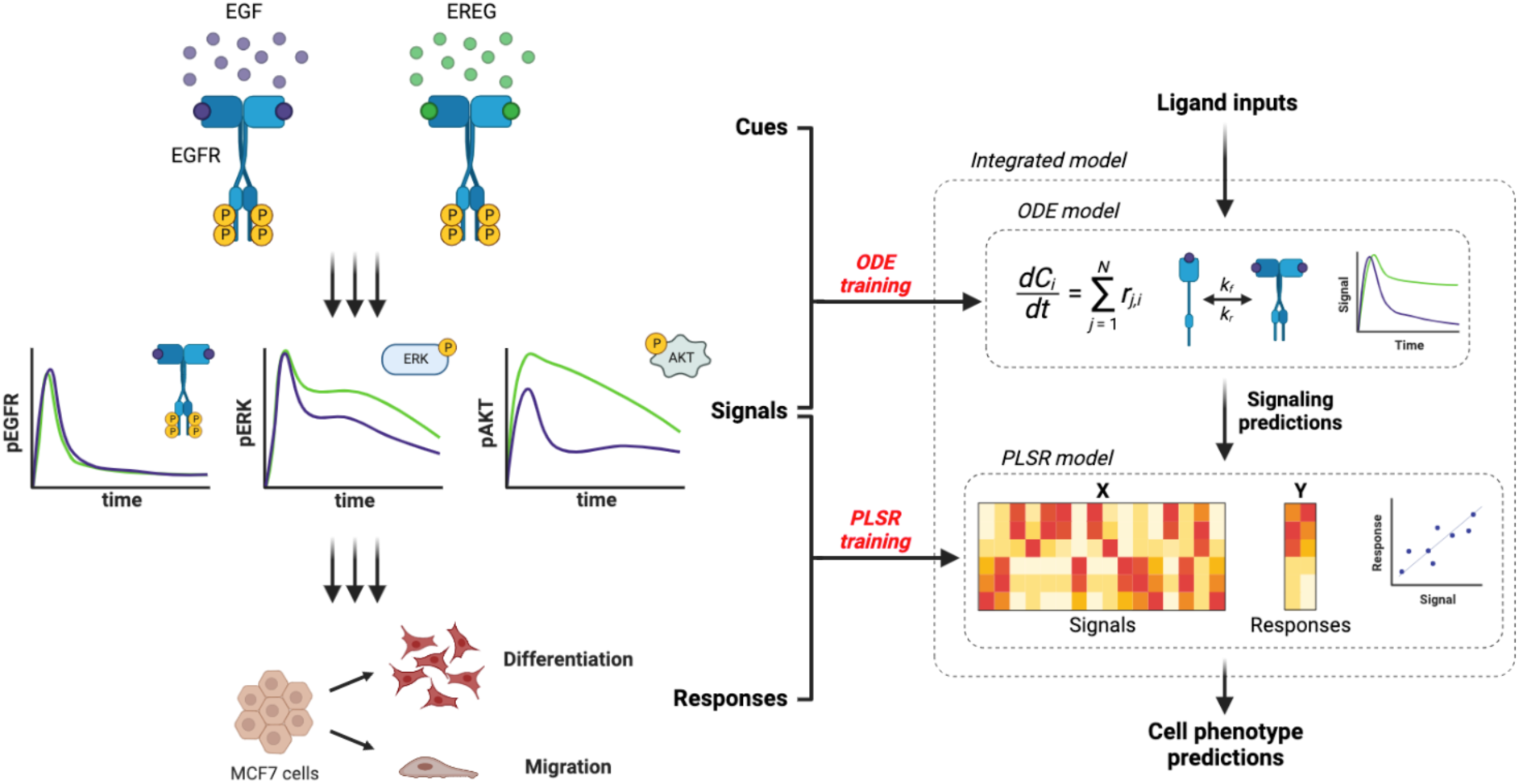
Integration of mechanistic and data-driven models enables prediction of phenotype dependencies on EGFR ligand properties. The integrated model consists of a mechanistic, ordinary differential equation (ODE) model describing the dynamics of the EGFR-ERK-AKT signaling network combined with a data-driven, partial least squares regression (PLSR) model that predicts MCF7 cellular responses to EGF and EREG (ligand inputs) using phosphoprotein predictions from the ODE model as inputs. The ODE and PLSR models are trained using dynamic phosphoprotein measurements of EGFR, ERK, and AKT, and the PLSR model is also trained on measurements of MCF7 migration and differentiation as responses.

**Figure 2.**
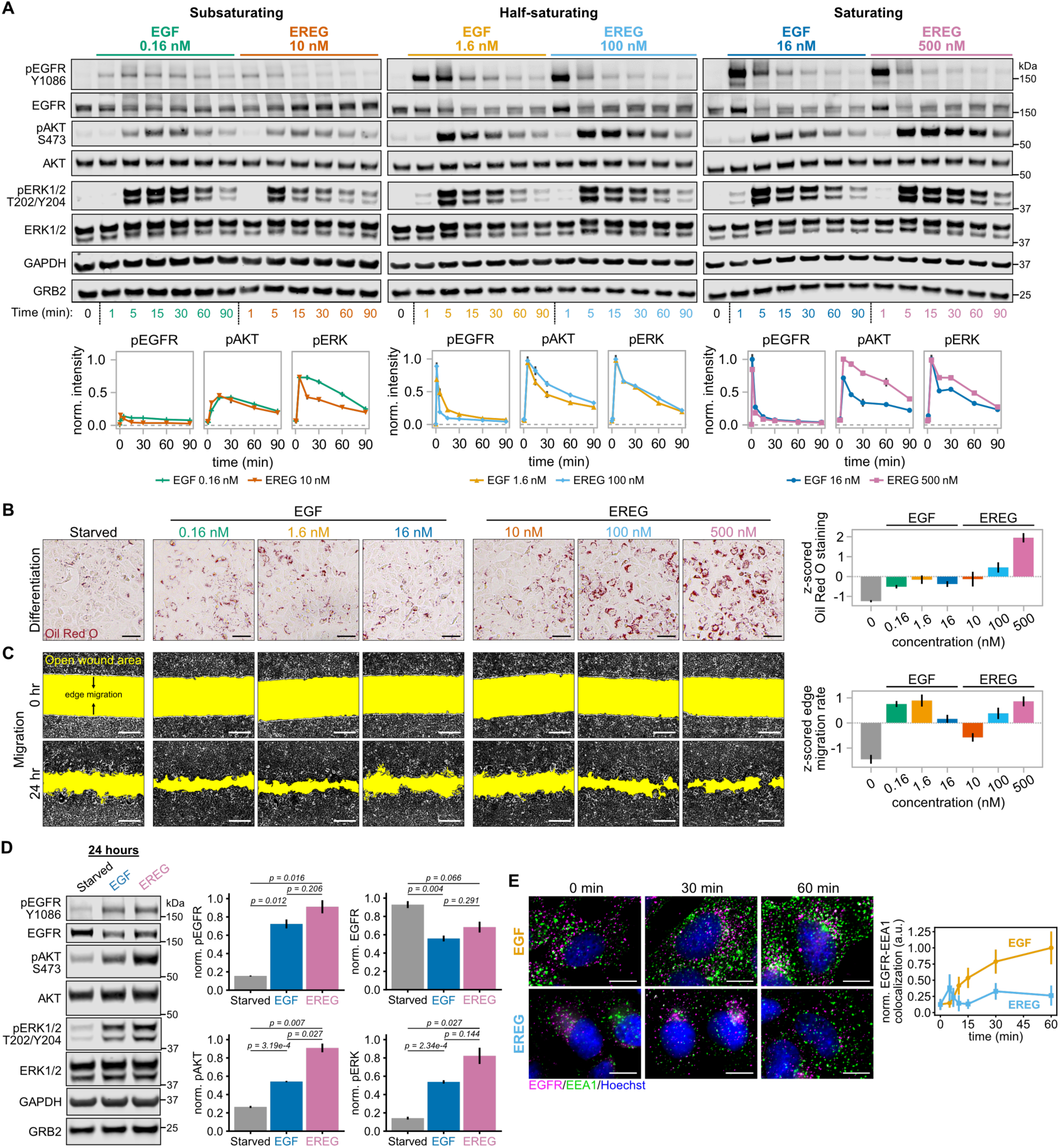
EGF and EREG induce differential signaling and cell phenotypes. **(A)** MCF7 cells were treated with EGF and EREG with the indicated concentrations and times, lysed, and analyzed by western blotting using antibodies against the indicated proteins. Phosphoprotein signals were first normalized by the summed intensities of GAPDH, GRB2, total ERK1/2, and total AKT, averaged across replicate membranes, and then normalized to the maximum value per analyte (norm. intensity). Error bars represent mean ± SEM, n = 3 blots. **(B)** Left: MCF7 cells were treated for 6 days with the indicated EGF and EREG concentrations or untreated (starved), fixed, and stained for neutral lipid accumulation using Oil Red O. Right: Oil Red O staining was quantified and z-scored (right). Error bars represent mean ± SEM, n = 3 biological replicates. Scale bars = 50 µm. **(C)** Left: MCF7 cells were treated and imaged for 24 hr for the same conditions as in **B** after removal of the two-well culture insert. The highlighted (yellow) area indicates the open wound area after removal of the silicone culture insert. Right: The mean rates of edge migration were calculated and z-scored. Error bars indicate mean ± SEM, n = 6 to 8 biological replicates per condition. Scale bars = 300 µm. **(D)** MCF7 cells were treated with EGF (16 nM), EREG (500 nM), or untreated (starved) for 24 hr, lysed, and analyzed by western blotting. Signals were normalized by the summed intensities of GAPDH, GRB2, ERK1/2, and AKT, averaged, and normalized to the maximum value across conditions. Error bars represent mean ± SEM, n = 3 blots. Statistical significance was determined using Welch’s one-way analysis of variance (ANOVA) and the Games-Howell post-hoc test. **(E)** Left: EGFR and EEA1 colocalization was analyzed by immunofluorescence in MCF7 cells treated with EGF (1.6 nM) or EREG (100 nM) for up to 60 min. Right: EGFR-EEA1 colocalization was quantified from immunofluorescence images as described in *Methods* and normalized by the maximum mean value across all conditions. Errors bars represent 95% confidence intervals, n = 64 to 442 cells per condition.

A potential explanation for differential downstream signaling dynamics between the ligands with modest or unapparent differences in receptor phosphorylation is that EREG induces slower EGFR internalization than EGF (*6, 45*). EGFR endocytosis attenuates AKT signaling by removing a driving force for membrane recruitment of PI3K, which phosphorylates the phosphoinositide PIP_2_ in the cascade leading to AKT activation (*25*). The effects of EGFR endocytosis on ERK1/2 activation are debated, with prior work identifying both positive and negative roles for endocytosis in regulating ERK signaling (*46–51*). In MCF7 cells, however, increased EGFR recycling and plasma membrane retention prolong ERK and AKT activation (*43, 44*). To confirm the anticipated trends in EGFR trafficking in MCF7 cells, we used immunofluorescence microscopy to demonstrate that EGFR colocalizes with the early endosome marker EEA1 significantly less in response to EREG than EGF (**Fig. 2E**). The long-term effect of this may underlie the larger residual total EGFR signal observed for EREG compared to EGF in Fig. 2D.

### Training the integrated model and initial model interpretations

As a first step in model training, the ODE model of EGFR-ERK-AKT signaling was fit to the data in Fig. 2A. The model was trained separately on the EGF and EREG datasets, which improved the quality of fitting compared to one universal fit. The need for two fits is sensible, given that EGF and EREG induce EGFR dimers with different structures and trafficking dynamics (*5, 6, 52*) and drive downstream signaling with different dynamics. Published mass spectrometry measurements in MCF7 cells (*53*) were used to estimate absolute phosphorylated ERK, AKT, and EGFR abundances (see *Methods*) (**Fig. 3A**). Fold-reduction in total EGFR by 90 min and total EGFR expression in MCF7 cells (*53*) were also used in fitting. The fitted model exhibited low residual error, with *R^2^* values of ∼0.92 and ∼0.99 for EGF and EREG data, respectively. The most notable model deviation from the data occurred for total EGFR, with the model under-predicting for the lowest doses of EGF and EREG by about 10-20%. Total EGFR levels, however, did not weigh heavily in the trained data-driven model, as described later. The mechanistic model was also tested by comparing predictions for endocytosed EGFR against experimental quantification of EEA1-EGFR colocalization for 1.6 nM EGF and 100 nM EREG, data which were not used in model training. While the amount of EREG-induced EGFR internalization predicted by the model is larger than observed experimentally, the model fit (which is not trained on receptor endocytosis data) does reflect that EREG induces less EGFR internalization than EGF at similar levels of receptor saturation, consistent with the experimental data (**Fig. 3B**). Similar model predictions are obtained when comparing the saturating doses for EGF and EREG (16 nM and 500 nM, respectively).

**Figure 3.**
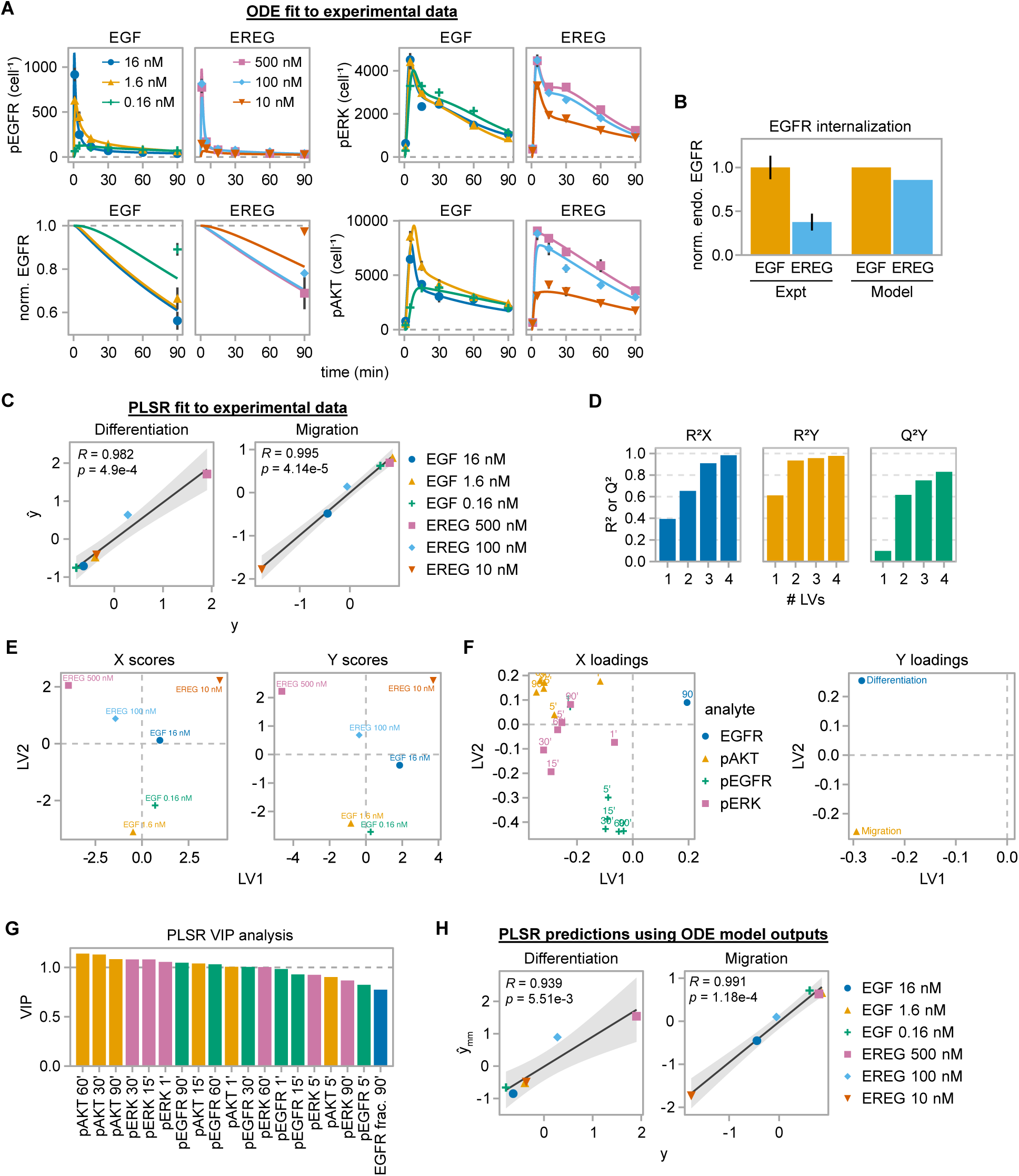
The integrated model accurately predicts EGFR-regulated phosphoprotein dynamics and cell phenotypes. **(A)** ODE model fits to experimental signaling dynamics were generated for the time courses of phospho-ERK, phospho-AKT, phospho-EGFR, and normalized EGFR expression (normalized to t = 0 min) for the indicated concentrations of EGF and EREG. **(B)** Normalized average levels of endocytosed EGFR (norm. endo. EGFR) were compared between experiments and model predictions. Quantification of EEA1-EGFR colocalization from Fig. 2E (Expt) and model predictions of endocytosed EGFR for up to 60 min of ligand treatment were time-averaged and compared for the 1.6 nM EGF and 100 nM EREG conditions. Data are normalized to the 1.6 nM EGF condition within each group. **(C)** Parity plots comparing PLSR model-predicted (ŷ) and scaled experimental (y) phenotype data are shown for the PLSR model trained on the data in Fig. 2, B and C. **(D)** PLSR model goodness of fit (R^2^X, R^2^Y) and prediction (Q^2^Y) metrics are plotted as a function of the number of latent variables. **(E)** X and Y PLSR scores are plotted for the first two latent variables. **(F)** X and Y PLSR loadings are plotted for the first two latent variables. Ligand treatment time (min) is indicated above each point for the X loadings. **(G)** Variable importance in projection (VIP) scores are shown for the trained PLSR model. **(H)** Comparisons between scaled experimental phenotype data (y) and PLSR predictions when using ODE model outputs as PLSR inputs (ŷ_mm_) are shown as parity plots. For **C** and **H**, the Pearson correlation coefficient and associated *p* value for the relationship between model predictions and experimental data are indicated on each plot. The best-fit linear regression line between model predictions and experimental data is shown as a black line, with 95% confidence intervals for the predicted relationship shaded in gray.

The data-driven model connecting signaling dynamics to phenotypes was based on PSLR, which is well-suited to scenarios with more features than independent conditions, as is the case here with 19 signaling measurement features for each of six ligand conditions. PLSR training used the signaling data from Fig. 2A and phenotype data from Fig. 2B-C as the independent (X) and dependent (Y) variables, respectively. Model quality was high, with goodness of fit R^2^X and R^2^Y ≈ 98% and goodness of prediction Q^2^Y ≈ 83% with four latent variables (LV) (**Fig. 3C-D**). Q^2^Y was calculated using leave-one-out cross-validation and indicates the model’s ability to predict held-out data when trained on the remaining data.

Separation of treatment conditions along LVs 1 and 2 in the scores plots reflects the model’s ability to discriminate among the ligand and concentration conditions used to generate the dataset (**Fig. 3E**). Comparison of those plots against the loadings (**Fig. 3F**) reflects the ability of the model to capture relationships among conditions, phenotypes, and signals. Phosphorylated ERK and AKT are highly loaded in LV1, in parallel with both phenotype loadings, but AKT projects positively with differentiation in LV2 while some, but not all, ERK measurements project negatively in LV2 with migration. While this suggests that ERK and AKT are both important regulators of differentiation and migration, the model predicts a tendency for AKT to control differentiation more strongly and for ERK to influence migration more strongly. EGFR phosphorylation projects strongly only into LV2, indicating that receptor phosphorylation dynamics do not determine phenotypes as much as downstream effectors. Variable importance in projection (VIP) analysis indicates that pAKT for *t* ≥ 30 min and pERK at 15 and 30 min were most explanatory of phenotypes (**Fig. 3G**). This is coincident with the largest variations in signaling occurring at those times, although this does not guarantee importance for predicting phenotypes in these types of models.

The trained models were coupled by using the ODE model predictions as inputs to the PLSR model, enabling predictions of phenotypic responses to EGF or EREG. Using ODE model predictions for the training conditions yielded accurate predictions of differentiation and migration responses in MCF7 cells (**Fig. 3H**), demonstrating that the integrated model performs well even with the inherent propagation of error from the ODE to the PLSR model.

### Model validation through pathway inhibition

We next validated the roles of ERK and AKT in determining ligand-induced MCF7 phenotypes by testing the effects of MEK and PI3K inhibition. Inhibitor effects on differentiation used 16 nM EGF and 500 nM EREG because these concentrations drove the strongest Oil Red O staining (**Fig. 2B**). Consistent with PLSR predictions and independent of the signaling and phenotype responses induced in the absence of inhibitors, the PI3K inhibitor GDC-0941 (PI3Ki) caused near-complete loss of Oil Red O staining for both ligands, whereas the MEK inhibitor trametinib (MEKi) had only modest or no effect (**Fig. 4A**). ERK1/2 and AKT1/2 knockdown experiments confirmed these roles for ERK and AKT in EGF- and EREG-induced MCF7 neutral lipid accumulation (**Fig. S1A-B**). The dominant role of PI3K/AKT in controlling differentiation was also observed with heregulin-β1, a HER3/ERBB3 ligand that drives strong AKT activation (**Fig. S1C**) (*5*). Thus, the relative importance of ERK and AKT for MCF7 differentiation is also independent of growth factor and receptor context. For testing the effects of inhibitors on migration, we used 0.16 nM EGF and 100 nM EREG, which are the lowest concentrations for each ligand that induce significant signaling and migration responses in MCF7 cells (**Fig. 2, A and C**). PI3K and MEK inhibition both substantially reduced cell migration in response to either ligand at these concentrations (**Fig. 4B**). Note that, in disagreement with our interpretation of Fig. 3F, the effect of PI3K inhibition was larger than that of MEK inhibition, a point we return to later.

**Figure 4.**
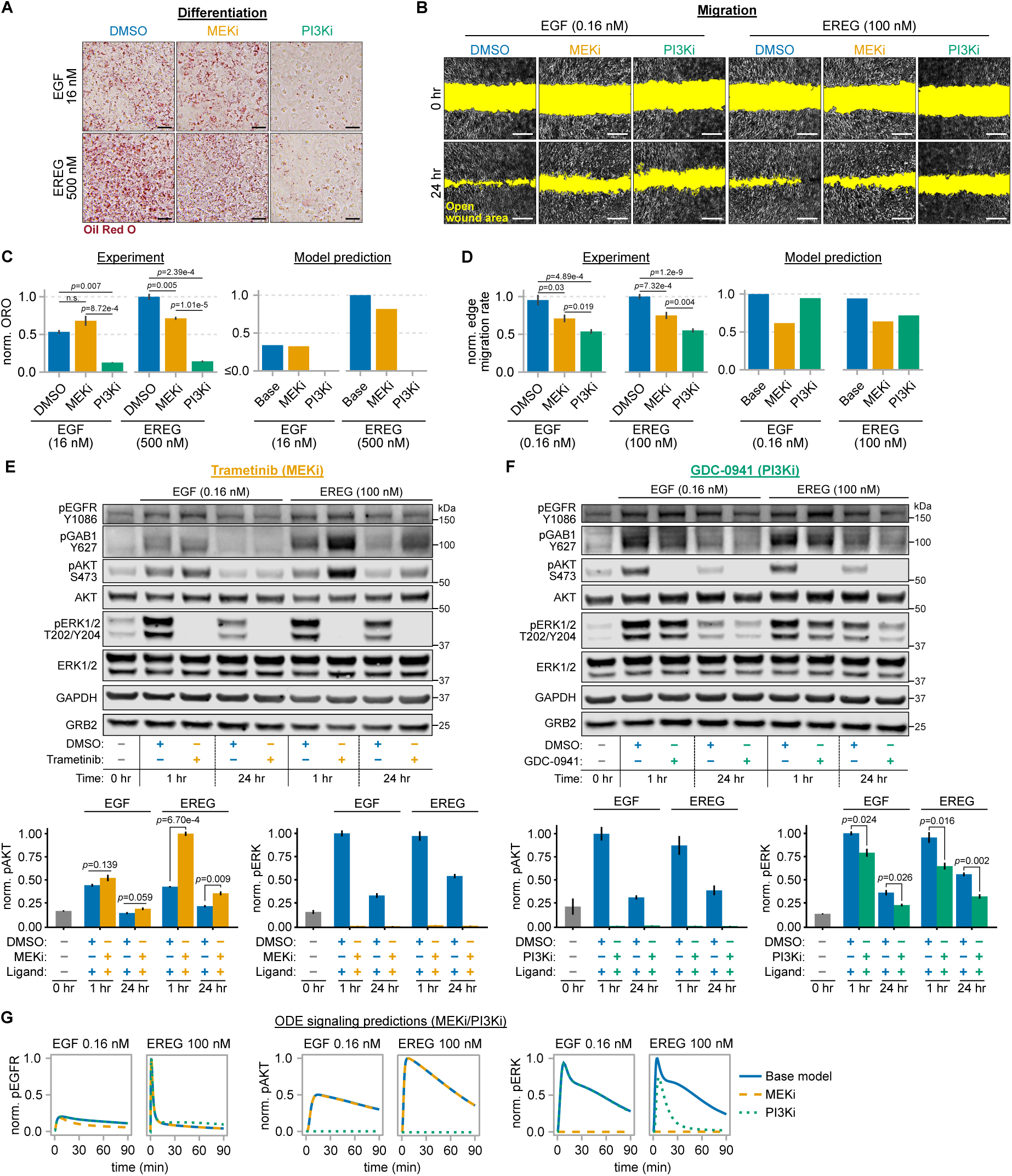
The integrated model predicts the phenotypic effects of MEK and PI3K inhibition. **(A)** MCF7 cells were treated with EGF (16 nM) or EREG (500 nM) and either DMSO, trametinib (MEKi; 50 nM), or GDC-0941 (PI3Ki; 50 nM) for 6 days and then stained with Oil Red O. **(B)** MCF7 cells were imaged for 24 hr after scratching and treatment with EGF (0.16 nM) or EREG (100 nM) and either DMSO, trametinib (50 nM), or GDC-0941 (50 nM). The highlighted (yellow) area indicates the open wound area after scratching. **(C)** Quantification of Oil Red O staining for the conditions in A (left) was compared against integrated model predictions (right) for simulated EGF or EREG treatments with or without MEK or PI3K inhibition. Data are normalized to the maximum value within the respective experiment or model prediction groups. To more directly compare to experimental data, model predictions were first transformed back to the original scale of the PLSR training data using the column standard deviations and means of the phenotype training data prior to maximum-value normalization. Error bars represent mean ± SEM, n = 4 biological replicates. Scale bar = 50 µm. **(D)** Mean edge migration rates quantified from **B** were compared against integrated model predictions for simulated EGF or EREG treatments for the base model or when simulating MEK or PI3K inhibition for the same EGF and EREG concentrations. Data are normalized to the maximum value within either the experiment or model prediction groups. Error bars represent mean ± SEM for at least 9 fields of view across 4 to 8 scratches per condition. Scale bar = 300 µm. **(E and F)** MCF7 cells were treated with EGF (0.16 nM) or EREG (100 nM) and either DMSO, trametinib (TRAM; 50 nM) **(E)**, or GDC-0941 (GDC; 50 nM) **(F)**, lysed, and analyzed by western blotting using antibodies against the indicated proteins. Phosphoprotein signals for ERK and AKT were normalized by the summed intensities of GAPDH, GRB2, total ERK1/2, and total AKT, averaged across replicate membranes, and then normalized to the maximum value per analyte (norm. intensity). Error bars represent mean ± SEM, n = 3 blots. **(G)** Model predictions for the dynamics of pEGFR, pAKT, and pERK for the base model and when simulating either MEK or PI3K inhibition. MEK and PI3K inhibition were simulated by setting the ODE model rate constants for ERK phosphorylation or PI3K-mediated PIP_2_ phosphorylation, respectively, to zero. Model predictions for each variable were normalized to the maximum value across simulations. Statistical significance in **C-F** was determined using Welch’s one-way analysis of variance (ANOVA) and the Games-Howell post-hoc test.

We next tested the ability of the integrated model to predict the quantitative effects of MEK and PI3K inhibition on phenotypic responses to ligands. Pathway inhibition was simulated by setting the rate constants for ERK phosphorylation by MEK and PIP_2_ phosphorylation by PI3K to zero. Using these ODE simulation results, the integrated model accurately predicts experimentally measured changes in differentiation in response to inhibitors (**Fig. 4C**), with a Pearson correlation coefficient of 0.951 (*p* = 0.003) between normalized experimental measurements and model predictions. The model is less accurate in predicting inhibitor effects on migration (**Fig. 4D**), with a Pearson correlation coefficient of 0.505 (*p* = 0.307). In particular, the model substantially under-predicts the effects of PI3K inhibition on migration, especially for EGF, which contrasts the original model prediction that pERK contributes more strongly to migration than pAKT. This is probably because pAKT is not weighted as heavily as pERK for predicting migration (**Fig. 3F**) and is not highly induced in response 0.16 nM EGF (**Fig. 2A**). Thus, for this concentration of EGF, the primary contribution to the model’s migration prediction comes from pERK, which is relatively high and sustained. While initial model calculations suggest that ERK may be the primary controller of MCF7 migration, these results indicate a significant and larger role for AKT in regulating migration. Continuing with the ligand concentrations used to test the effects of MEKi/PI3Ki on migration, Western blotting confirmed that ERK and AKT phosphorylation were durably suppressed by inhibitors of the respective pathways (**Fig. 4E-F**). Interestingly, however, AKT phosphorylation was significantly elevated in response to EREG+MEKi. This coincided with increased phosphorylation of GAB1, the adaptor that localizes PI3K to EGFR and the cell membrane and thus enables PI3K to phosphorylate PIP_2_ (*54*). This effect may occur due to antagonism of ERK-mediated GAB1 phosphorylation on multiple serine and threonine residues (T312, S381, S454, T476, S581, S597) that inhibits GAB1-PI3K binding (*55*), an effect not captured in the ODE model. The preferential occurrence of this effect for EREG may be due to the slow internalization of EREG-EGFR signaling complexes, which promotes AKT activation. In response to PI3K inhibition, AKT phosphorylation was suppressed in ligand-treated cells for at least 24 hr (**Fig. 4F**). ERK phosphorylation was also slightly suppressed by PI3K inhibition. This effect was smaller in EGF-treated cells but statistically significant for both ligands at 1 and 24 hr. The decrease in ERK phosphorylation may be linked with the suppressed phosphorylation of GAB1 Y627, which engages N-terminal SH2 domains on the protein tyrosine phosphatase SHP2 to promote complete ERK activation downstream of many receptor tyrosine kinases (*56, 57*).

Model-predicted pEGFR, pAKT, and pERK dynamics for MEK and PI3K inhibition were assessed to confirm the accuracy of the predicted signaling effects on which phenotypic predictions are based. For MEK or PI3K inhibition, the model predicted little or no effect on EGFR phosphorylation (**Fig. 4G**), consistent with experiments. The model also predicted no effect of MEK inhibition on AKT phosphorylation, consistent with EGF but not EREG experiments. This is not surprising, given the lack of model accounting for effects such as the ERK-GAB1 feedback (*55*) previously mentioned. For PI3K inhibition, the model predicted a substantial decrease in ERK phosphorylation for EREG only and failed to predict the observed effect in the presence of EGF (**Fig. 4G**). This occurs because the model predicts significant EREG-induced GAB1-PIP_3_ binding, which provides a route for GAB1-bound GRB2-SOS membrane localization and thus SOS-catalyzed RAS activation. While there is no direct evidence for this mechanism, the ability of membrane-targeted SOS to efficiently activate RAS (*58*) and the attenuation of ERK activation when GAB1 membrane localization is antagonized by overexpression of PTEN or GAB1’s PH domain (*15*) suggest that it is possible. The lack of PI3K inhibition effect on predicted EGF-induced ERK phosphorylation also explains the predicted minor effect of PI3K inhibition on EGF-induced migration (**Fig. 4B**).

The failure to predict increased AKT phosphorylation in response to trametinib led us to ask if the model would still accurately predict inhibitor-dependent phenotypes if model-predicted pAKT levels were adjusted to match the measurements in Fig. 4E. This is effectively a test of the PLSR model’s ability to predict phenotypes given corrected signaling data. When the AKT dephosphorylation rate constant was decreased independently for EGF and EREG to reproduce the observed changes in pAKT observed with MEK inhibition (**Fig. 5A**), the integrated model accurately predicted the small increase in Oil Red O staining for EGF+MEKi but erroneously predicted a large increase in Oil Red O staining for EREG+MEKi (**Fig. 5B**). This points to a failure of the PLSR model, trained using ligand response data alone, to predict phenotypes when the signaling features (independent variables) are perturbed using inhibitors. Because PLSR uses a linear combination of weighted independent variables, any amount of ERK inhibition can be compensated for by sufficient AKT phosphorylation. However, the observation that some level of ERK activity is required for complete MCF7 differentiation (**Fig. 4A**) and the results in Fig. 5B suggest that an interaction exists between ERK and AKT that is missing in the PLSR model.

**Figure 5.**
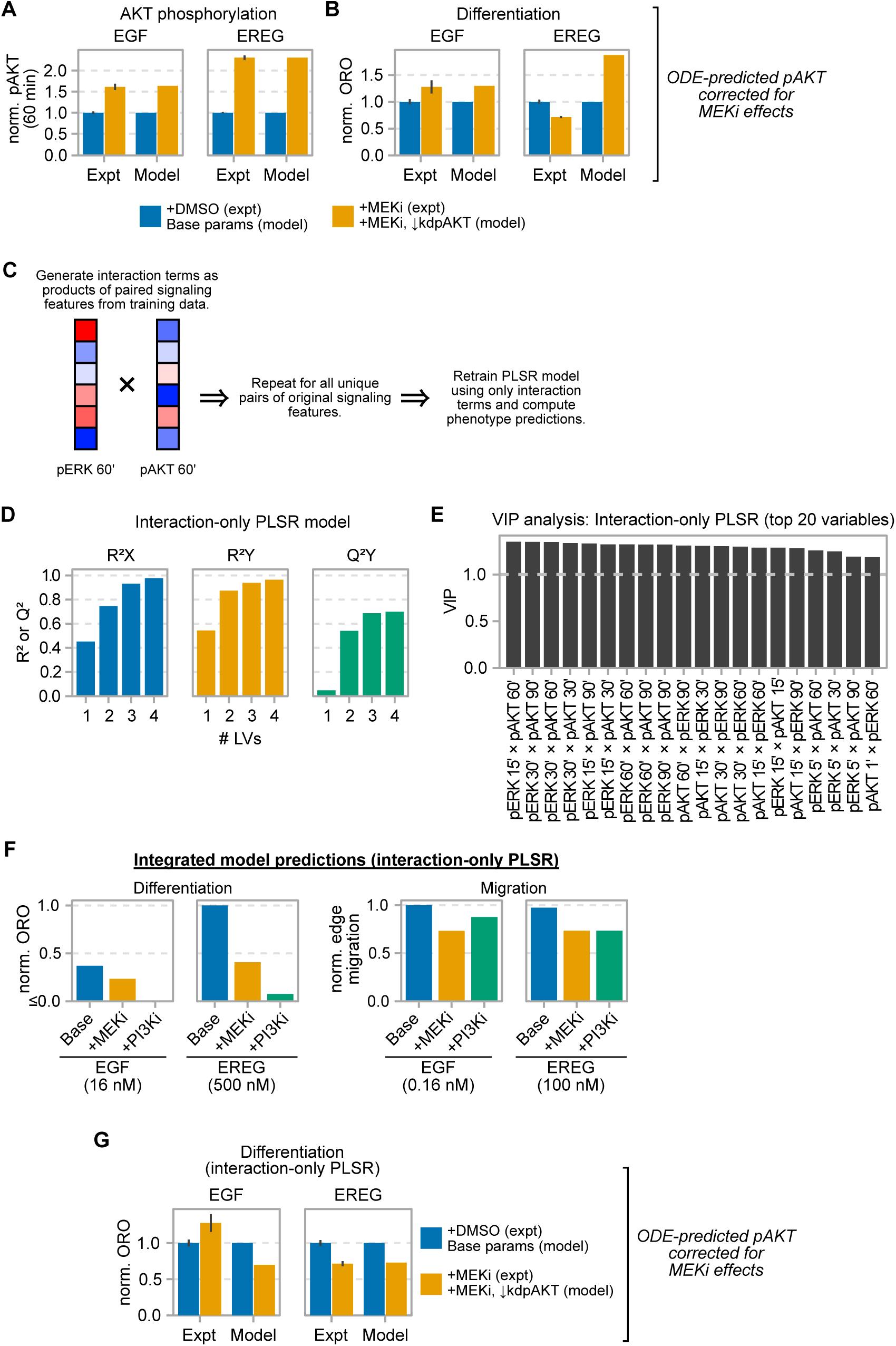
Incorporating interaction terms in the PLSR model improves predictions of inhibitor-induced phenotypes. **(A)** Experimental pAKT quantification (Expt) at 60 min of growth factor treatment from Fig. 4E was compared against mechanistic model pAKT predictions for 0.16 nM EGF and 100 nM EREG with or without trametinib (+MEKi) at 60 min when decreasing the AKT dephosphorylation rate constant (*kdpAKT*) to match the experimentally observed ratio between the DMSO and trametinib conditions for each ligand. Model predictions are normalized to the maximum value for each ligand. Within each pair of bars, data are normalized to the +DMSO/Base model condition. **(B)** Experimental Oil Red O quantification (Expt) from Fig. 4A was compared against integrated model predictions of Oil Red O staining for 16 nM EGF and 500 nM EREG were computed when using the values of *kdpAKT* from **A** to generate mechanistic model predictions and were compared against experimental Oil Red O quantification (Expt) for the same conditions. Within each pair of bars, data are normalized to the +DMSO/Base model condition. **(C)** The PLSR model was retrained using only interaction terms, which were computed as the products between all unique pairs of signaling variables from the original PLSR training data. **(D)** Model goodness of fit (R^2^X, R^2^Y) and prediction (Q^2^Y) metrics are plotted as a function of the number of latent variables for the interaction-only PLSR model. **(E)** The top 20 variable importance in projection (VIP) scores are shown for the interaction-only PLSR model. **(F)** Integrated model predictions for Oil Red O staining or the rate of edge migration were computed for the indicated conditions using the interaction-only PLSR model. Predictions are normalized to the maximum value within the experiment or model prediction groups. **(G)** Experimental Oil Red O quantification (Expt) from Fig. 4A was compared against integrated model predictions of Oil Red O staining for 16 nM EGF and 500 nM EREG, which were computed with the interaction-only PLSR model when using the values of *kdpAKT* calculated in **A** to generate mechanistic model predictions. Within each pair of bars, data are normalized to the +DMSO/Base model condition. Model predictions in **B**, **D**, and **F** were first transformed back to the original scale of the PLSR training data using the column standard deviations and means of the phenotype training data prior to normalization.

To incorporate ERK-AKT interactions, we generated a revised PLSR model trained only on interaction variables computed as pairwise products of every unique combination of timepoints for distinct analytes (**Fig. 5C**). The resulting “interaction-only” PLSR model fit the training data well (R^2^Y ≈ 98%) and exhibited good performance in cross validation (Q^2^Y ≈ 83%) with four latent variables (**Fig. 5D**). VIP analysis indicates that interactions between later-time (≥ 30 min) pERK and pAKT variables are most predictive of phenotypes and that interactions involving pEGFR were of only minor importance (**Fig. 5E**). For simulations of MEK and PI3K inhibition, this interaction-only version of the model produced slightly improved predictions compared to the original non-interaction model (**Fig. 5F)**, but it still underpredicts the importance of AKT for migration relative to ERK, in part because other variables not involving AKT can still contribute significantly to migration predictions. However, when correcting the predicted effects of MEK inhibition by adjusting the AKT dephosphorylation rate to reproduce the observed increases in AKT phosphorylation in the ODE model predictions, the updated interaction-only model correctly predicts the decrease in Oil Red O staining observed with EREG+MEKi (**Fig. 5G**; compare to Fig 5B). The Oil Red O staining prediction for EGF+MEKi, however, was less accurate for this version of the model. A potential explanation is that some of the excluded non-interaction terms from the original PLSR model are required to predict the effects of EGF+MEKi on differentiation.

### Model sensitivity analysis

The unique structure of the integrated model enables calculation of phenotype sensitivities to changes in model parameters. The extended Fourier amplitude sensitivity test (eFAST) was first used to compute multivariate parameter sensitivities where ODE model initial protein concentrations were simultaneously varied by factors ≤100 from assumed or best-fit values. Calculations were performed for 20 log-linearly spaced EGF or EREG concentrations to assess concentration dependence. For EGF, differentiation and migration predictions were most sensitive to the abundance of EGFR, followed by AKT, PP2A, and GAB1 (**Fig. 6A**), and differentiation predictions were relatively sensitive to PDK1, PI3K, and PIP_2_ perturbations. Sensitivities were generally invariant with changes in EGF concentration (**Fig. 6B**). Given the importance of AKT in determining phenotypes, it is not surprising that many top sensitivities, especially for differentiation, are associated with AKT signaling. The relatively high sensitivity to PP2A changes occurs because PP2A dephosphorylates AKT, RAF, and MEK, and thus represents a single node for regulation of both phenotype-determining pathways. Although VIP analysis did not nominate pEGFR as a substantial controller of phenotypes, the high eFAST sensitivity to total EGFR expression indicates that the receptor plays an important role in determining phenotypes by setting the absolute activation levels of downstream effectors. This effect is particularly apparent for migration, which is substantially more sensitive to EGFR expression changes than to changes in other signaling nodes. Migration predictions are particularly sensitive to EGFR because perturbing EGFR expression simultaneously perturbs downstream AKT and ERK activation, both of which are important determinants of model migration predictions (**Fig. 3F**). Similar sensitivity results are seen for EREG (**Fig. 6C-D**), with the most notable difference being the higher first-order sensitivities (eFAST main effects) for EGFR, which indicates a dominant control of model outputs by variations in EGFR expression. The greater sensitivity of EREG migration calculations compared to EGF is consistent with the observation that AKT and ERK activation are more sensitive to changes in the concentration of EREG compared to EGF across similar levels of receptor saturation (**Fig. 3A**). In general, however, the total sensitivities of model predictions (main effects plus interactions) to changes in EGFR expression versus other proteins were largely consistent between the ligands for both phenotypes, with minor exceptions being the lower sensitivity of EREG-induced migration to PP2A and PDK1 (**Fig. 6, A and C**). Additionally, model phenotype predictions for both ligands were sensitive to at least one adaptor molecule (GAB1 for EGF, GRB2 for EREG), consistent with observations that the abundance of adaptor proteins is a critical determinant of EGFR-initiated signaling (*53*). Overall, the small magnitudes for most main effect indices for both ligands indicate that phenotype predictions are dictated largely by interactions among multiple species rather than by one pathway node alone.

**Figure 6.**
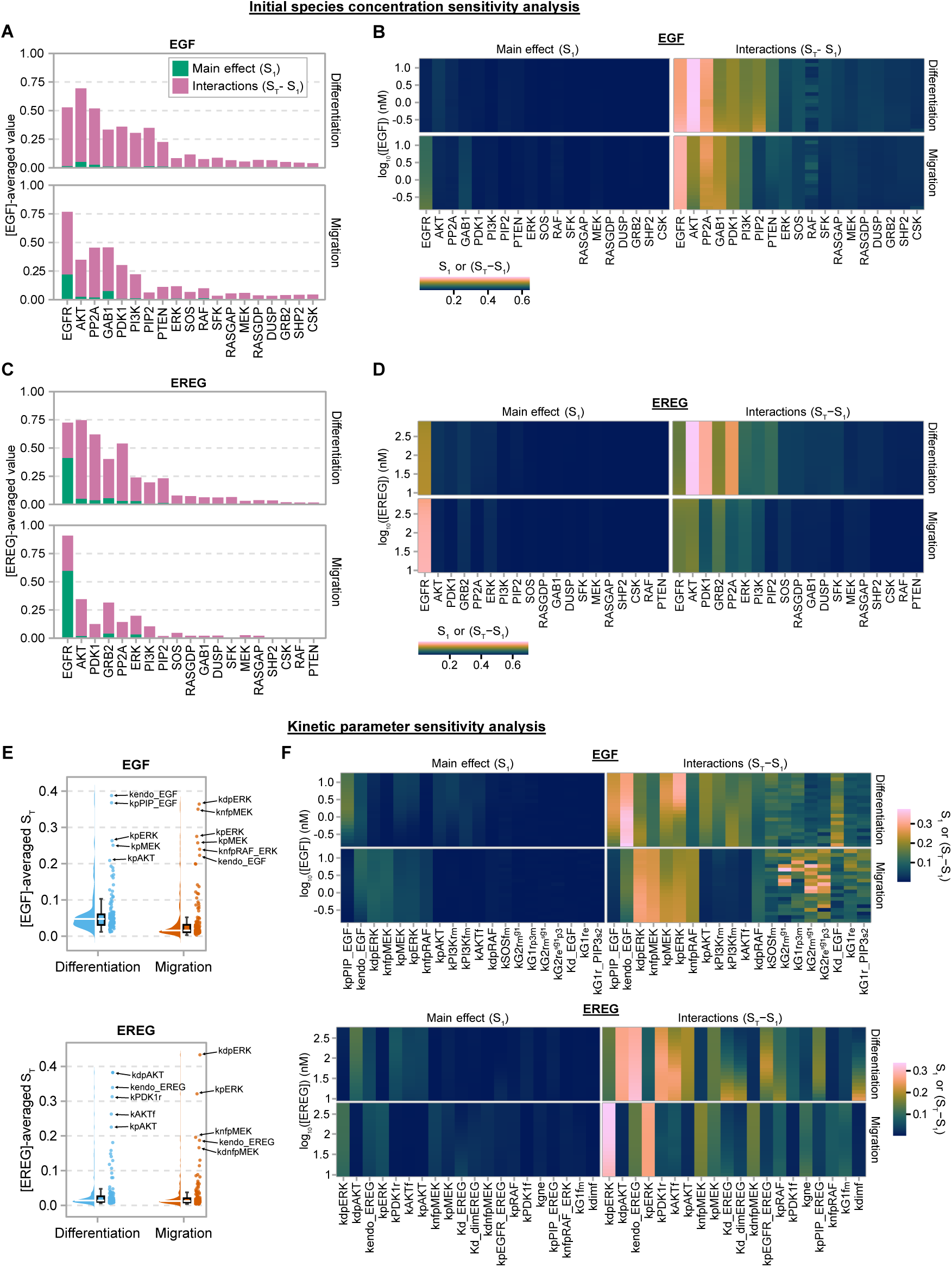
Global sensitivity analysis identifies proteins and kinetic processes determining model phenotype predictions. **(A)** Extended Fourier amplitude sensitivity test (eFAST) was used to calculate sensitivities for integrated model phenotype predictions for EGF treatment when varying the initial concentrations of the indicated ODE model species over 20 log_10_-linearly spaced EGF concentrations between 0.16 and 16 nM. First- and total-order eFAST indices (S_1_ and S_T_, respectively) were computed for each concentration, followed by calculating the interaction index as S_T_−S_1_. The concentration-averaged indices were calculated by integrating with respect to log_10_-transformed EGF concentrations. Parameters are ranked by their summed first-order and interaction indices across the two phenotypes. **(B)** The concentration-dependent eFAST indices used in the calculations in **A** are plotted in the heatmap as functions of log_10_-transformed EGF concentration. **(C)** Concentration-averaged model eFAST indices for EREG treatment were calculated as in **A**. **(D)** The concentration-dependent eFAST indices used in the calculations in **C** are plotted as a heatmap as functions of log_10_-transformed EREG concentration. **(E)** Total-order eFAST indices were computed and averaged for integrated model phenotype predictions when varying ODE model kinetic parameters for each of the concentrations of EGF and EREG in **B** and **D**, respectively. **(F)** The concentration-dependent eFAST indices used in the calculations in **E** are plotted as functions of log_10_-transformed EGF or EREG concentration for the top 20 most-sensitive parameters for each ligand, as determined by summing the concentration-integrated S_T_ values across both phenotypes for each parameter.

eFAST was also applied to the ODE model kinetic parameters to identify key reactions and interactions. Based on the distributions of the total-order eFAST indices, which indicate the total variance in model output explained by a parameter’s main effect and interactions with other parameters, phenotype predictions for EGF and EREG were highly sensitive to only a small number of parameters (**Fig. 6E**). For both ligands, most parameters explained < 10-15% of model output variance. Model output variance was largely explained by parameter interactions, as indicated by relatively large interaction indices (**Fig. 6F**). Consistent with the analysis in Fig. 6A-D, parameters related to EGFR (e.g., *kendo_EGF/EREG*, *Kd_EGF/EREG*, *Kd_dimEREG*, and *kdimf*) and AKT activation (e.g., *kdpAKT*, *kPDK1f/r*, *kpPIP_EGF/EREG*, *kPI3Kf/r*, etc.) were among the most phenotype-controlling for both ligands (**Fig. 6F**). Model predictions were also highly sensitive to parameters related to ERK activation and deactivation (*kpERK*, *kdpERK*), consistent with the fact that ERK is still an important regulator of cell phenotypes, particularly migration. For both ligands, the appearance of several parameters describing the activation of proteins immediately upstream of ERK (i.e., RAS, RAF, and MEK) and AKT (i.e., PI3K and PDK1) suggests the importance of interactions and feedback mechanisms that regulate the activity of these upstream effectors in determining phenotypes. Multiple parameters related to GRB2 and GAB1 binding are also among the top sensitive parameters for EGF, and one parameter describing to GAB1-GRB2 binding, *kG1f_m_*, was among the top 20 most sensitive parameters for EREG, again highlighting adaptor proteins as contributing to signaling and phenotypic outcomes (**Fig. 6F**).

A notable observation is that the rate constants describing EGFR internalization (*kendo_EGF/EREG*) were the second- and third-most sensitive parameters for EGF and EREG, respectively (**Fig. 6F**). Many studies have demonstrated the importance of receptor internalization and trafficking, particularly of EGFR, in determining the strength and duration of receptor-mediated signaling and phenotype responses (*6, 12, 24, 43*). Our experiments show that EGF and EREG induce distinct EGFR internalization dynamics (**Fig. 2E**), and the integrated model’s sensitivity to the rate constants for EGFR internalization (**Fig. 6F**) suggests that differential EGFR trafficking is one of the key mechanisms by which EGF and EREG elicit distinct signaling and phenotypes, in addition to differential effects on EGFR activation and dimerization (*5*). Model predictions are also highly sensitive to perturbations in the rate constants and abundances of species involved in processes and feedbacks acting on EGFR-downstream effectors (e.g., *kdpERK*, *knfpRAF*, *knfpMEK*, *kdpAKT*). Overall, model calculations here suggest that cellular responses to different EGFR ligands are determined by integrated effects at multiple levels of the signal transduction cascade that are differentially activated by high- or low-affinity ligands.

## DISCUSSION

Here, we demonstrate how the integration of mechanistic and data-driven models of signaling, in this case applied to EGFR, can be used to overcome a critical gap in our understanding – how quantitative perturbations at the signaling input level propagate through the earliest mechanistic steps in signal initiation and through multivariate downstream signaling pathways to effect changes in cell phenotypes. A key to accomplishing this type of model integration lies at the interface of the two modeling approaches, the signaling output nodes of the mechanistic model that serve as inputs to the data-driven model. On one hand, those signaling outputs must be limited in scope to avoid the attendant explosion of parameters that occurs as nodes are added to differential equation-based models. At the same time, there must be enough relevant signaling pathway outputs to predict phenotypic outcomes accurately. Striking that balance will be critical and challenging in new applications of this approach, but it may ultimately enable our understanding of one of the most challenging questions in receptor signaling biology, namely how the same receptors can drive alternative phenotypes in different cell contexts (*59, 60*).

The ability of different EGFR ligands to specify alternative cell outcomes is typically attributed to regulation of the ERK pathway (*5, 7, 9, 43*). Studies comparing EGFR to other receptors have also focused on ERK as the phenotype-controlling pathway, as in the classical example of PC12 cells treated with EGF or nerve growth factor to drive preferential proliferation or differentiation, respectively (*61*). The integrated modeling approach we developed here correctly predicted that ERK and AKT cooperate to drive differentiation and migration in MCF7 cells and that AKT plays the dominant role for differentiation. The model structure enabled us to pinpoint ligand-mediated receptor trafficking as a controlling rate process that defines phenotype specification for high- and low-affinity EGFR ligands. At the receptor level, the phenotypic impacts of different EGFR ligands have separately been attributed to effects on receptor dimerization and activation kinetics (*5, 52, 62, 63*) and ligand-dependent receptor trafficking and degradation (*43, 64–66*). Of course, all these processes are coupled. Because dimerization strength directly affects the lifetime of active EGFR, ligand-specific differences in EGFR degradation and internalization are inextricably coupled to dimerization dynamics. This is reflected in phenotype predictions that are largely determined by interactions of EGFR dimerization and internalization parameters with other model parameters (**Fig. 6F**).

The integrated model structure allowed us to assess the effects of protein expression on phenotype predictions. While model predictions for both ligands are highly sensitive to EGFR expression, they are also generally sensitive to the expression of downstream adaptors and phosphatases (**Fig. 6, A and C**). The primary exception is that EREG migration predictions are notably less sensitive to the expression of proteins other than EGFR due to the greater sensitivity of ERK and AKT activation to changes in the concentration of EREG compared to EGF (**Fig. 3A**). Because the expression of some EGFR pathway members can vary by orders of magnitude across cell contexts and in some cases represent stoichiometric bottlenecks for signal transduction(*53*), it is not surprising that model predictions are also sensitive to the expression of EGFR-downstream pathway members. For example, EGF predictions are sensitive to the concentration of GAB1, which is expressed at low copy numbers (∼2,800 molecules/cell) compared to its downstream binding partners SHP2 and PI3K (∼54,000 and ∼200,000 molecules/cell, respectively)(*53*). Other lowly expressed proteins relative to downstream effectors in MCF7s include EGFR (∼2,500 molecules/cell), SOS (∼4,900 molecules/cell), and RAF (∼40,000 molecules/cell), though only EGFR ranked highly in the model sensitivity analysis. Collectively, our results suggest that EGFR-initiated signaling dynamics and phenotype outcomes are regulated by integrated inputs of nodes and kinetic processes across multiple levels of the signaling network that can be influenced by the binding properties of the EGFR ligand.

The improvement in model predictions observed when using features based on products of signaling measurements suggests that pathways cooperate to determine some phenotypes (e.g., in the presence of pathway inhibitors). The implied cooperativity could involve ERK and AKT regulating the same targets or different targets that drive the same phenotypes. As an example of the former model, the ERK and AKT pathways share some transcription factor targets (e.g., ATF/CREB, MEF2, SRF, and PBX1) (*67*) that are important regulators of migration and proliferation responses (*68–71*). As an example of the latter model, AKT, but not ERK, promotes lipogenesis by upregulating SREBP family transcription factor activity (*72*), but sustained cytoplasmic ERK activity promotes the stability of mRNAs necessary for MCF7 differentiation (*44*). Thus, one hypothesis is that ERK stabilizes AKT-regulated mRNAs that promote neutral lipid accumulation, and this may involve ERK and AKT activities that occur asynchronously in time. This possibility is captured in the model by the interaction variables involving ERK and AKT phosphorylation time points. Even in the classical example of PC12 cells, evidence for pathway cooperativity can be found. In single-cell immunofluorescence imaging of NGF- or EGF-treated PC12 cells, simultaneous consideration of phosphorylated ERK and AKT better predicts PC12 differentiation versus proliferation than does consideration of ERK alone (*73*). A counter-example is the observation that SK-N-BE(2) neuroblastoma cell differentiation in response to EREG stimulation is primarily ERK-dependent, as simultaneous EREG treatment with MEK inhibitors but not PI3K inhibitors significantly reduces neurite outgrowth (*9*). Thus, the pathways controlling phenotypes such as differentiation can be cell line- and context-dependent. The ability to incorporate cell-line specific protein expression as part of the mechanistic model means that the integrated modeling framework described here can be readily applied to other cell backgrounds to investigate the context dependence of cell phenotypes, though this may require retraining the model with data from the cell line(s) of interest.

For incorporating crosstalk among signaling variables, multiple approaches for generating model features are available. Kernel ridge regression and support vector machines can perform automatic variable transformation and feature generation using polynomials, the radial basis function, and other kernel functions. For models not based on kernels, manual feature generation is possible. Features involving, for example, ratios or products of two or more variables, each with a unique exponent, can be used. In gene-phenotype models of bacterial antibiotic resistance, incorporating features describing pairwise and three-way interactions among genes significantly improves the ability of specialized LASSO regression models to predict the effects of antibiotics on bacterial proliferation (*74*). However, because the number of new features that can be generated is essentially limitless, rigorous cross-validation and feature selection should be used to tune models with large numbers of generated features to avoid overfitting (*75*). A promising option to circumvent manually specifying interactions among predictor variables is the use of data-driven models such as CellBox, which automatically infers network connectivity during model training (*76*). Network inference approaches generally require data for a large number of pharmacological perturbations, but this can be aided by use of publicly available data repositories such as the Cancer Perturbed Protein Atlas, which contains time-resolved phosphoproteomic and drug response data (*40*).

The present model only considers ERK and AKT as downstream effectors, but EGFR activates other pathways that regulate migration and proliferation, including PLCγ (*77–79*) and STAT1/3/5 (*80, 81*). The model can be updated to incorporate or substitute these pathways as necessary to predict phenotypes where those pathways are controlling, with the previously mentioned caveat that adding pathways expands the number of model parameters. Additionally, consideration of subcellular localization and cytosolic concentration gradients of signaling effectors is important for accurately describing signal transduction and cellular decision-making (*12, 51, 82-84*). Incorporating spatial or compartmental information about signaling effectors (e.g., distinct variables for membrane-localized, cytoplasmic, or nuclear pERK) may therefore be another avenue to improve model performance. Training such models is substantially more complicated, however, as it requires measurements of signaling effectors with subcellular resolution and the potential use of partial differential equations. Thus, the computational cost of training such models could become prohibitive.

## METHODS

### Computational Methods

#### Mechanistic model formulation

The mechanistic, ordinary differential equation (ODE) model was implemented in the Julia programming language (*85*) (version 1.8.3). The model was constructed by specifying chemical reactions describing the EGFR signaling network using Catalyst.jl (*86*). Reactions are assumed to follow mass-action kinetics by default. Model reactions were compiled as a system of ODEs using DifferentialEquations.jl (*87*) and solved with the TRBDF2 solver (*88, 89*), which performs well for highly stiff systems. Simulation efficiency was aided by symbolically computing the Jacobian of the ODE system using ModelingToolkit.jl (*90*).

The topology is based on prior EGFR signaling models (*13, 20, 27, 29, 30*) and describes the EGFR-initiated activation of the RAS/ERK and PI3K/AKT signaling axes. In total, the model includes 757 reactions (including reversible and irreversible steps), 127 species, and 149 parameters. The distinct proteins described by the model include EGFR, GRB2, SOS, GAB1, SHP2, RAS, RAF, MEK, ERK, PI3K, AKT, PTEN, PP2A, PIP_2_, PDK1, dual-specificity phosphatases (DUSPs 1, 4, and 6, lumped into a single species), Src family kinases (SFKs), C-terminal Src kinase (CSK), and RAS p21 protein activator 1 (RASGAP/RASA1). Expression of AKT, PTEN, PI3K, SFKs, CSK, PP2A, and DUSP1 was estimated by scaling the absolute numbers of these proteins in HeLa cells (*91*) based on the ratio of mRNA transcript abundances (transcripts per million) for the corresponding genes in MCF7 and HeLa cells from the Cancer Dependency Map (*92*). Total DUSP concentration was calculated by summing the estimated expression of DUSP1 and absolute expression levels of DUSP4 and DUSP6 estimated directly from mass spectrometry (*91*). PIP_2_ was assumed to be expressed at ∼8,000 molecules/μm^2^ on the plasma membrane (*93*), or ∼1×10^7^ molecules/cell. For the remaining species, expression was specified as previously described (*53*). The concentrations of initially non-zero model species are provided in **Table S1**. A brief overview of the signaling processes included in the model is described below.

Signaling in the model is initiated by binding of EGF or EREG to cell-surface EGFR. Ligand-bound EGFR undergoes dimerization and autophosphorylation, which we assume occurs on a single representative tyrosine residue. We similarly assume that other model species, such as GAB1, are phosphorylated on a single residue to simplify the model topology and avoid the combinatorial complexity introduced by considering multiple phosphorylation sites (*94, 95*), which places a severe computational burden on model simulations and parameter estimation. Phosphorylated EGFR serves as a binding site for the adaptor GRB2, which we assume can simultaneously bind the guanine exchange factor SOS and the adaptor protein GAB1, each to one of GRB2’s two SH3 domains. The regulation of SFK activity is complex and involves separate phosphorylation and dephosphorylation steps for the positive regulatory tyrosine (Y418 on Src) and the C-terminal negative regulatory tyrosine (Y530 on Src) (*96, 97*). As in a previous model (*29*), SFK activation was approximated as a process catalyzed by phosphorylated EGFR, and SFK deactivation was approximated as a process catalyzed by CSK, the enzyme responsible for phosphorylating Src on Y530 (*97*). GAB1 phosphorylation was modeled as a process catalyzed by either active SFKs (*29*) or by EGFR when GAB1 is bound to it (*98, 99*). Because PI3K and SHP2 bind to GAB1 on the distinct phospho-tyrosine residues 472 (*100*) and 627 (*101, 102*), respectively, we assume GAB1 can simultaneously bind free SHP2 and PI3K molecules once phosphorylated in the model. We assume that GAB1 dephosphorylation occurs as a zeroth-order process with respect to phosphatases, as in a previous model (*29*). Because SH2 domain binding protects phospho-tyrosines from phosphatase activity (*103, 104*), we assume GAB1 can only be dephosphorylated when neither SHP2 nor PI3K are bound to it.

Membrane-localized SOS catalyzes the activation of RAS, which catalyzes the sequential activation of the RAF-MEK-ERK cascade. Based on the ability of membrane-targeted SOS to efficiently activate RAS (*58*) and the observation that ERK kinase activity is attenuated when GAB1 membrane localization is antagonized by overexpression of PTEN or GAB1’s PH domain (*15*), we allow for the possibility that RAS can be activated by SOS bound to GRB2:GAB1:PIP_3_ complexes at the membrane. Importantly, we assume that RAS exists only on the plasma membrane and thus cannot be directly activated by EGFR-bound SOS on endosomes based on data that suggest RAS exists primarily on the plasma membrane (*51, 105*). Membrane-localized PI3K phosphorylates PIP_2_ to PIP_3_, which serves as a binding site for cytosolic PDK1, AKT, and GAB1-containing protein complexes. AKT can be phosphorylated once bound to PDK1:PIP_3_. The direct binding of GAB1-containing complexes to PIP_3_ (*27*) provides another route for SOS and PI3K membrane localization and thus activation and phosphorylation of RAS and PIP_2_, respectively. We also allow for any PIP_3_-bound GAB1 signaling complex to decouple from PIP_3_ and bind directly to EGFR, either through binding of GAB1 to EGFR-bound GRB2 or GRB2 binding to phosphorylated EGFR.

Feedbacks described in the model include EGFR degradation in endosomes, ERK-mediated phosphorylation of SOS (*106*) and of RAF and MEK (*14*), and RASGAP-catalyzed conversion of RAS-GTP to RAS-GDP (*107*). Any GRB2-bound EGFR species can undergo internalization to endosomes, where EGFR can either be recycled back to the membrane or degraded. As in previous models, we include separate rates and recycling fractions for ligand-bound and -unbound receptors on endosomes (*13, 108*). Adaptor binding to EGFR continues on endosomes, and we assume that EGFR degradation causes immediate unbinding of any receptor-bound adaptor complexes. Active, GTP-bound RAS can be deactivated through direct binding of RASGAP, which catalyzes the conversion of RAS to its inactive, GDP-bound state. Unbinding of RASGAP from RAS can be promoted by membrane-localized SHP2 (through binding to GAB1). Multiple ERK-mediated feedbacks are included in the model, including ERK phosphorylation of SOS, RAF, and MEK (*14, 106*). We assume that feedback phosphorylation of SOS causes its immediate unbinding from GRB2 and that feedback phosphorylation of RAF and MEK inhibits their ability to phosphorylate and activate MEK and ERK, respectively.

#### Mechanistic model parameter estimation

To aid in parameter estimation and accommodate the possibility that significantly different model parameter values may be required to adequately fit the data for each ligand, we generated separate mechanistic model fits against EGF and EREG signaling data. In total, 124 of the 149 model parameters were fit to experimental data. The model was fitted to estimates for the total number of phosphorylated of EGFR, ERK, and AKT and to the estimated fraction of EGFR expression after 90 min of ligand treatment (the final time point), all of which were calculated using dynamic phosphoprotein data gathered from western blotting (**Fig. 2A**). Western blot data for phospho-EGFR (pEGFR), phospho-ERK (pERK), and phospho-AKT (pAKT) were first averaged across replicate membranes and normalized to the maximum value for each analyte across all EGF and EREG treatment conditions at 1, 5, 15, 30, 60, and 90 min of ligand treatment. Estimates for absolute numbers of pERK, pAKT, and pEGFR molecules were then calculated from the normalized western blot data based on prior mass spectrometry estimates of absolute expression of EGFR-MAPK pathway members and of phosphorylated ERK1/2 and EGFR in response to EGF (*53*). Based on these data, we estimate that ∼2% of ERK1/2 and ∼10% of EGFR are phosphorylated at maximum in MCF7 cells. This fraction of EGFR phosphorylation is consistent with the other cell lines studied by Shi et al. (*53*), and the fraction of phosphorylated ERK in MCF7s, while significantly lower than the ∼30% phosphorylated in the other tested cell lines, is consistent with the significantly lower expression levels of EGFR in MCF7s (∼2,500 receptors per cells) compared to other cell lines in the same study (>150,000 receptors per cell). The maximum fractional level of AKT phosphorylation was assumed to match that of ERK phosphorylation. The fraction of total EGFR at 90 min was calculated for each ligand condition from western blot data by dividing the normalized EGFR intensity at 90 min of ligand treatment by the normalized EGFR intensity for untreated cells. These estimates were then used to scale the normalized western blot data to obtain estimates for absolute numbers of phosphorylated proteins for fitting and constraining the mechanistic model.

Parameter fitting was performed by minimizing the sum of squared errors between model outputs and the experimental signaling data described above. Because the scales of the fitting variables varied significantly due to using absolute numbers of molecules, the fitting errors were normalized by the experimental values so that all data points for all variables contributed relatively equally to the fitting process. To further aid the fitting process, we manually performed iterative optimization runs on the data, first beginning with data from the earlier timepoints and sequentially adding later time points, using the best-fit parameter values from the prior optimization run as initial guesses for fitting the data with the next set of time points. Model fitting was first completed for the full set of data for the highest and lowest concentrations of each ligand (16 nM and 0.16 nM for EGF; 500 nM and 10 nM for EREG), followed by adding the data for the intermediate concentrations (1.6 nM for EGF; 100 nM for EREG). Minimization of the sum of squared errors was performed using the evolutionary centers algorithm (*109*) from the Metaheuristics.jl library (*110*). To allow the model to more easily fit the training data, we fit most model parameter values using initial guesses and nominal values taken from prior models or experimental measurements (see **Table S2**). Model parameters describing EGF and EREG binding to EGFR and EGFR dimerization were pinned during fitting so that model predictions would be constrained based on known EGFR binding and dimerization properties for these ligands (described below). We also pinned rate constants describing GTP hydrolysis by RAS and RAS-bound RASGAPs (i.e., conversion of active RAS-GTP to inactive RAS-GDP) based on experimental measurements for RAS- and RASGAP-catalyzed hydrolysis (*107*). EGFR recycling fractions (ligand-bound or - unbound) were allowed to vary between 1.0 and 1×10^-12^. All other model parameter values were allowed to vary by up to six factors of 10 above or below their initial values during fitting.

EGF and EREG binding to EGFR and EGFR dimerization were modeled as reversible processes occurring at the plasma membrane and within endosomes with forward binding rate constants estimated assuming diffusion-limited binding (*111*). Average forward binding rate constants for EGF and EREG binding to EGFR were calculated by first computing forward binding rate constant curves for each ligand on the plasma membrane and on endosomes using the range of possible free EGFRs (1 to 2,486 receptors per cell) (*53*) and the surface areas of the cell and endosomes (*111*). Cells and endosomes were approximated as spheres of radii 10 μm and 350 nm (*112*), respectively, and the diffusivities of EGF and EREG were both assumed to be 1.5×10^-6^ cm^2^/min (*113*). The resulting rate constant curves were then integrated to obtain averaged receptor-ligand forward binding rate constants for each ligand on the plasma membrane and endosomes. Ligand-receptor reverse binding rate constants were calculated using the forward binding constants and an effective binding affinity of EGF for EGFR of 1.6 nM (*114*). The effective EREG binding affinity for EGFR was estimated to be ∼76 nM by scaling the effective EGF-EGFR binding affinity of 1.6 nM by the ratio of EGF and EREG binding affinities with sEGFR501 (16 nM and 760 nM, respectively) as measured by isothermal calorimetry (*5*). Average forward rate constants for EGFR dimerization were similarly estimated assuming diffusional limitations (*111*) and an EGFR diffusivity of 2.16 µm^2^/min.

#### PLSR modeling

Partial least squares regression (PLSR) was performed using the *pls* R package (*115*). To integrate PLSR calculations with ODE model simulations, the *pls* package was called within Julia using RCall.jl (https://github.com/JuliaInterop/RCall.jl). Averaged western blot (signaling) data and phenotype data were mean-centered and variance-scaled column-wise prior to model training. Column means and scales (variances) were recorded and used to rescale ODE model outputs prior to inputting them to the PLSR model. The optimal number of PLSR latent variables was selected using leave-one-out cross-validation by determining the number of latent variables that maximized the “goodness of prediction” metric Q^2^Y (i.e., minimized prediction error on held-out data). For the interaction-term PLSR model, new predictor variables were generated from the original set of signaling training data by computing the products between all possible pairs of variables prior to mean-centering and variance-scaling.

#### Mechanistic and PLSR model integration

The mechanistic and PLSR models were integrated to predict MCF7 phenotype responses to EGFR ligand binding by first generating mechanistic model predictions for a given dose of EGF or EREG using the best-fit parameter values for the ligand of interest. The mechanistic model predictions corresponding to the predictor variables of the trained PLSR model were then extracted from the ODE outputs and centered and scaled using the recorded column means and variances for the PLSR training (X) data. The centered and scaled mechanistic model predictions were then used as inputs to the PLSR model to obtain integrated model predictions of MCF7 differentiation and migration responses for the original EGF or EREG dose of interest.

#### Integrated model parameter sensitivity analysis

The sensitivity of integrated model phenotype predictions to simultaneous changes in mechanistic model initial species abundances or kinetic parameters was assessed using the extended Fourier amplitude sensitivity test (eFAST) (*116, 117*). eFAST calculates first-order (S_1_) and total-order sensitivity indices (S_T_) for each input parameter by simultaneously varying all parameters between manually specified upper and lower bounds (see below) according to sine waves with distinct frequencies and using Fourier decomposition to calculate the variance in model output explained by the input parameters. S_1_, also known as the main-effect index, accounts for the fraction of overall model output variance that can be attributed solely to an individual parameter, whereas S_T_ indicates the fraction of the overall model output variance that can be attributed to a parameter both individually and when accounting for all possible interactions with the other model parameters being varied. Thus, the variance in model output that can be attributed solely to interactions of an input parameter with other model parameters is calculated as the difference S_T_–S_1_. Separate eFAST analyses were performed for initial species concentrations and for kinetic parameters for both EGF and EREG. Kinetic parameters and initial protein concentrations were allowed to vary by up to a factor of 100 above or below their base values. The best-fit values for kinetic parameters were used as their base values. In all eFAST calculations, 1,000 simulations were run per varied model parameter.

### Experimental Methods

#### Cell culture

MCF7 cells (ATCC, HTB-22) were cultured in Dulbecco’s Modified Eagle’s Medium (DMEM) supplemented with 10% fetal bovine serum (FBS) (Avantor), 1% penicillin-streptomycin (10,000 µg/mL), and 1% L-glutamine (200 mM) in a humidified incubator at 37°C and 5% CO_2_. Cells were serum starved for at least 16 hr in DMEM supplemented with 0.5% FBS, 1% penicillin-streptomycin, and 1% L-glutamine prior to treatment with growth factors or inhibitors. Cells were confirmed to be mycoplasma-negative using the MycoAlert Mycoplasma Detection Kit (LT07-318, Lonza), and the identity of MCF7 cells was confirmed by STR profiling through the Johns Hopkins University Genetic Resources Core Facility cell line authentication service.

#### Growth factors and inhibitors

Recombinant human EGF (PeproTech, AF-100-15), EREG (PeproTech, 100-04), and heregulin-β1 (PeproTech, 100-03) were diluted in starvation medium prior to addition to cells. The MEK inhibitor trametinib (ApexBio, A3018x) was used at 50 nM, and the PI3K inhibitor GDC-0941 (Cayman Chemical, 11600) was used at 50 nM. Stocks of both inhibitors were prepared in DMSO. Cells were pre-incubated with inhibitors for 30 min prior to addition of growth factors.

#### Oil Red O staining

MCF7 cells were plated in 24- or 96-well tissue culture plates at a density of 1×10^5^ cells/cm^2^ (2×10^5^ or 3.2×10^4^ cells per well, respectively). ∼24 hr after plating, cells were serum starved and then treated with growth factors and/or inhibitors for 6 days without media changes. Cells were then fixed in 4% paraformaldehyde for 20 min, incubated in 60% isopropanol for 5 min, incubated with Oil Red O staining solution for 5 min to stain neutral lipids, and then washed at least 3 times with phosphate buffered saline (PBS). Oil Red O staining solution was prepared by dissolving Oil Red O powder (Alfa Aesar, A12989) at a concentration of ∼1.14 mg/mL in 60% isopropanol, followed by filtering the resulting solution through a 0.22 µm membrane (Millipore, SCGPT05RE). Hoechst DNA stain was added at a dilution of 1:2000 just prior to staining cells with Oil Red O. After Oil Red O staining and PBS washing, cells were imaged in-well in PBS on a BioTek Cytation 5 at 20× magnification using DAPI and Cy5 filter cubes for Hoechst and Oil Red O, respectively. Fluorescence images of Oil Red O and Hoechst stains were background-corrected, deconvolved, thresholded, and segmented using BioTek’s Gen5 software. Cell numbers were identified based on the segmented nuclear (Hoechst) stain, and Oil Red O staining was quantified as the area of thresholded Oil Red O fluorescence divided by the number of identified cells. The same fluorescence threshold was applied for individual channels across all images. Representative images of Oil Red O staining were also obtained using color brightfield at 20× magnification.

#### Wound healing assays

Wound healing assays were performed using either removable two-well silicone culture inserts (ibidi, 80209) or a standard scratch assay, as noted in figure legends. For culture insert-based wound healing assays, the inserts were placed in individual wells of 24-well tissue culture plates, and MCF7 cells were then seeded onto the culture area enclosed by the inserts at a density of ∼175,000 cells/cm^2^ (∼38,500 cells per chamber of the culture insert). For scratch-based assays, cells were seeded onto bare 24-well culture plates at the same density (∼350,000 cells per well). 24 hr later, cells were switched to starvation medium for at least 16 hr prior to removal of the culture insert or scratching with a 200 µL micropipette tip. After insert removal or scratching, cells were briefly washed at least twice with warm starvation medium, followed subsequently by treatment with growth factors and/or inhibitors and imaging on a BioTek Cytation 5 with onstage incubation at 37°C and 5% CO_2_. Cells were imaged at 10× magnification with phase contrast every 15 min for up to 24 hr. For wound healing experiments involving inhibitors, cells were pretreated with inhibitors for at least 30 min prior to scratching and washing steps. For all wound healing experiments, images were background-subtracted and smoothed using Gen5 software and then analyzed with a custom CellProfiler (*118*) pipeline to identify the area of the cell-free gap in each image. Briefly, images exported from Gen5 were illumination-corrected, inverted, histogram-equalized, and then smoothed with a Gaussian filter using a typical artifact diameter of 25 pixels. Smoothed images were thresholded to identify cell-free gaps as primary objects. The average distance between the gap edges (edge migration distance, EMD) was calculated as previously described (*43*) using the area of the cell-free gap and known image dimensions. The average rate of edge migration was calculated by numerically estimating the derivative of each EMD curve using finite differences, followed by taking the mean value of the resulting derivatives.

#### Western blotting

Cells were lysed using a standard lysis buffer (Thermo Fisher, FNN0011) supplemented with protease and phosphatase inhibitors (Sigma-Aldrich, P8340, P5726, P0044) and PMSF (Thermo Fisher, 36978). Crude lysates were centrifuged for 10 min at 15,000 rpm at 4°C, and the resulting supernatants were removed as clarified lysates. Lysate protein concentrations were determined with a micro-bicinchoninic acid (BCA) assay (Pierce, 23225). Approximately 30 µg of lysate were combined with 10× (500 mM) dithiothreitol (DTT) reducing agent, NuPage 4X LDS sample buffer (Invitrogen, NP0007), and high-purity water to reach equal sample volumes.

Samples were then heated at 100°C for 10 min and loaded onto a 1.5 mm NuPAGE 4-12% gradient gel (Invitrogen, NP0336BOX). After electrophoresis, proteins were transferred to 0.2 μm nitrocellulose membranes using the Trans-Blot Turbo Transfer System (Bio-Rad) using a 15 V constant voltage and 1.3 A maximum current for 30 min. Membranes were blocked with 4% bovine serum albumin (BSA) (Gemini Bio, 700-109P) dissolved in Tris-buffered saline (Bio-Rad #1706435) supplemented with 0.025% Triton X-100 (TBST). Primary antibodies were diluted in Intercept Blocking Buffer (IBB) (LI-COR, 927-60001) or in TBST (see below) and were incubated on membranes overnight at room temperature. Membranes were then rinsed three times with water, washed for 3 min with water, and then washed twice for 3 min with TBST. Infrared dye-conjugated or horseradish peroxidase-conjugated secondary antibodies were diluted 1:10,000 in TBST and incubated on membranes with shaking for 2 hr at room temperature. Membranes were then washed with water and TBST as before. For near-infrared (NIR) immunofluorescence detection, membranes were imaged on a LI-COR Odyssey CLx. For enhanced chemiluminescence (ECL) detection, membranes were incubated for 5 min in SuperSignal West Pico PLUS chemiluminescent substrate (Thermo Fisher, 34579) according to manufacturer’s instruction just prior to detection on a Bio-Rad ChemiDoc. Infrared dye-conjugated secondary antibodies were purchased from Rockland Immunochemicals (anti-mouse IgG DyLight 680 conjugated, 610-144-121; anti-rabbit IgG DyLight800 conjugated, 611-145-002). Horseradish peroxidase-conjugated secondary antibodies were purchased from Promega (anti-rabbit IgG, W4011). Membranes were stripped with 0.2 M NaOH as needed, with confirmation of stripping by re-imaging. Image Studio software (LI-COR) was used to quantify band intensities. The following primary antibodies were used with NIR detection at 1:1000 in IBB: AKT pS473 (CST, #4060), AKT (CST, #9272), ERK1/2 pT202/pY204 (CST, #4370), ERK1/2 (CST, #4695), GAPDH (Santa Cruz Biotechnology, sc-32233), GRB2 (Santa Cruz Biotechnology, sc-255). The following primary antibodies were used with ECL detection and were diluted in TBST: EGFR pY1086 (CST, #2220, 1:500), EGFR (CST #4267, 1:500) and GAB1 pY627 (CST, #3233, 1:1000).

#### siRNA-mediated ERK1/2 knockdown

siRNAs against ERK1/2 (#6560) and AKT1/2 (#6211) and SignalSilence control siRNA (#6568) were purchased from Cell Signaling Technology and were used at 10 µM. MCF7 cells were plated at a density of 375,000 cells/cm^2^ and simultaneously transfected with siRNAs using Lipofectamine RNAiMAX (Thermo Fisher) using the reverse transfection protocol, per manufacturer guidelines. Cells were seeded at twice the density used for assessing neutral lipid accumulation to account for cell death caused by the transfection process. Cells were then subjected to serum starvation and growth factor treatment 24 hr after transfection. 6 days after initial growth factor treatment, cells were fixed in 4% paraformaldehyde and stained with Oil Red O as described above. Cells were stained by indirect immunofluorescence by permeabilizing in 0.25% Triton X-100 in PBS for 5 min, followed by incubation with anti-ERK1/2 (CST 4695, 1:800) or anti-AKT (CST 9272, 1:200) primary antibody overnight at 4°C. Following five washes with 0.1% Tween 20 in PBS, cells were incubated in-well with Alexa Fluor Plus 488 anti-rabbit secondary antibody (Thermo Fisher, A32790) and Hoechst nuclear stain (1:2,000) for 1 hr at 37°C. Stained cells were imaged in-plate on a BioTek Cytation 5. Oil Red O staining and ERK1/2 fluorescence intensity were quantified from images using BioTek’s Gen5 software. All antibodies were diluted in IBB.

#### Coverslip immunofluorescence of EGFR-EEA1 colocalization

MCF7 cells were plated on 18-mm glass coverslips at a density of ∼3,800 cells/cm^2^ in 10 cm plates and allowed to adhere for 24 hr. Cells were then starved and treated with growth factors for up to 60 min. After growth factor treatment, cells were placed on ice, washed with ice-cold PBS twice, and fixed in 4% paraformaldehyde for 20 min. Fixed cells were washed twice with PBS and permeabilized with 0.25% Triton-X in PBS. Permeabilized cells were then stained overnight at 4°C with primary antibodies against EGFR (Santa Cruz, sc-120, 1:400), EEA1 (CST #3288, 1:400), and ERK 1/2 (Alexa Fluor 546-conjugated; Santa Cruz sc-514302, 1:200) diluted in IBB. Following five 3-min washes with 0.1% Tween 20 in PBS, coverslips were incubated with Alexa Fluor 488 anti-rabbit and Alexa Fluor 594 anti-mouse secondary antibodies (Thermo Fisher, A11034 and A11005, respectively) and Hoechst nuclear stain (1:2000) for 1 hr at 37°C. Coverslips were washed again with 0.1% Tween 20 in PBS, mounted on glass slides using Prolong Gold Antifade Mountant (Thermo Fisher, P36930), and sealed with clear nail polish. Z-stacks of fluorescence images were acquired on a Zeiss Axiovert Observer.Z1 fluorescence microscope using a 63× oil-immersion objective lens. At least eight fields of view and at least 100 cells were imaged per coverslip. Deconvolved, maximum-intensity *z*-projections were imported into CellProfiler and analyzed for colocalization of EGFR and EEA1 signals. Images were corrected for uneven illumination, and background-subtracted using Laplacian-of-Gaussian and top-hat filters. Individual nuclei were identified as primary objects by thresholding the Hoechst nuclear stain, and cell bodies were identified as secondary objects by thresholding the ERK1/2 stain. EGFR and EEA1 primary objects were identified from thresholded images. EGFR objects were classified based on overlap with EEA1 objects, and EGFR-EEA1 colocalization was quantified on a per-cell basis by computing the mean fluorescence intensity of EEA1-overlapping EGFR objects and dividing by the mean fluorescence intensity of all EGFR objects within each cell.

---

**Supplemental Figure S1.**
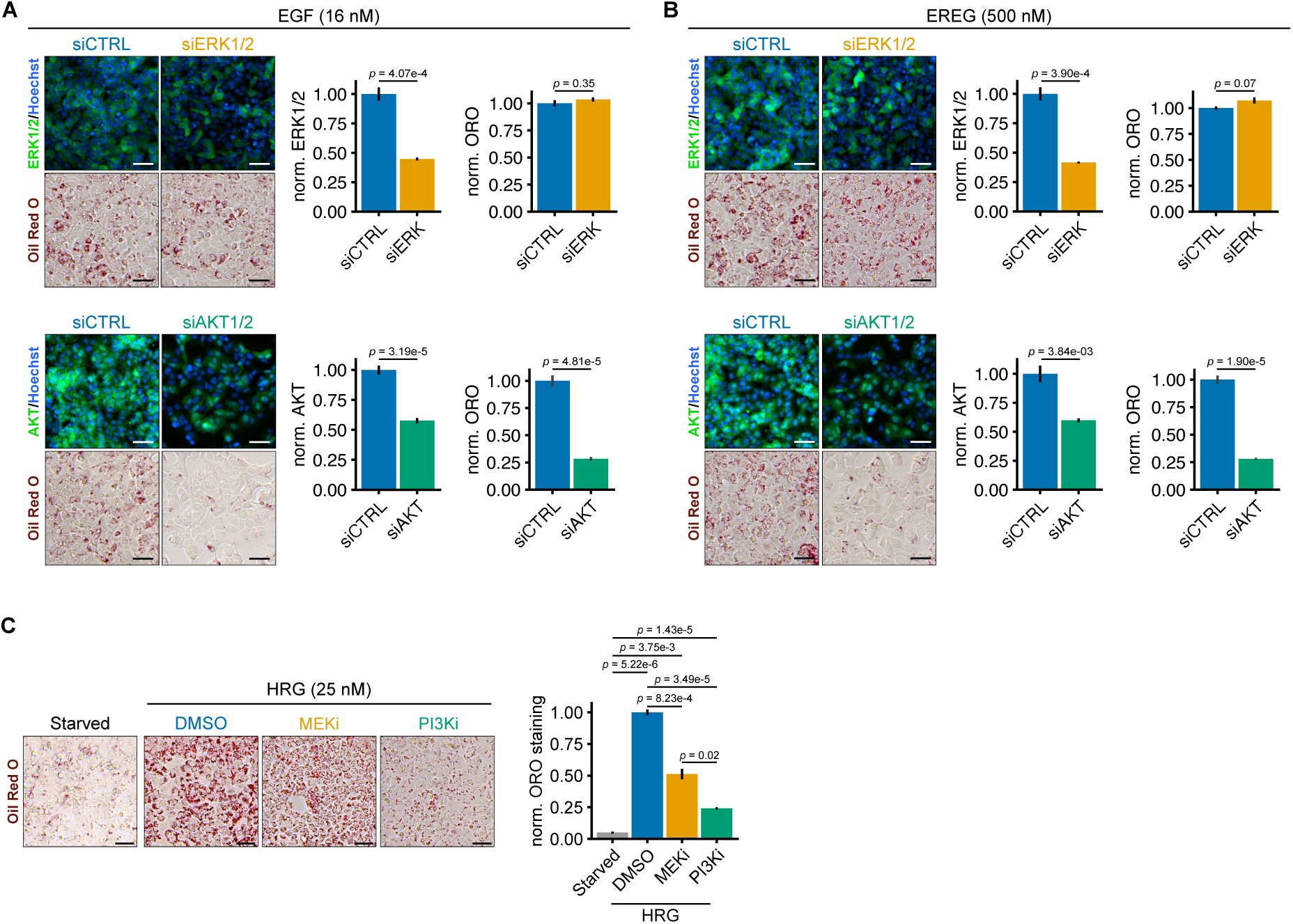
PI3K inhibition and AKT1/2 knockdown more strongly antagonize ErbB-mediated MCF7 differentiation than MEK inhibition and ERK1/2 knockdown. **(A)** Left: MCF7 cells were transfected with control, ERK1/2-targeting, or AKT1/2-targeting siRNAs, treated for 6 days with EGF (16 nM), and stained with Oil Red O and an antibody for ERK1/2 or AKT as indicated. Right: Oil Red O staining and integrated ERK1/2 or AKT intensity was quantified and normalized to the maximum values for each variable (right). Error bars represent mean ± SEM, n = 5 biological replicates. **(B)** Left: MCF7 cells were transfected with control, ERK1/2-targeting, or AKT1/2-targeting siRNAs, treated for 6 days with EREG (500 nM), and stained with Oil Red O and an antibody for ERK1/2 or AKT as indicated. Right: Oil Red O staining and integrated ERK1/2 or AKT intensity was quantified and normalized to the maximum values for each variable (right). Error bars represent mean ± SEM, n = 5 biological replicates. Statistical significance in panels **A** and **B** was assessed using Welch’s t-test. Scale bar = 50 µm. **(C)** Left: MCF7 cells were untreated (starved) or treated with heregulin-β1 (HRG; 25 nM) plus DMSO, trametinib (50 nM), or GDC-0941 (50 nM) for 6 days and then stained with Oil Red O. Right: Oil Red O staining was quantified and normalized to the maximum value. Error bars represent mean ± SEM, n = 4 biological replicates. Statistical significance was determined using one-way ANOVA with the Games-Howell post-hoc test. Scale bar = 50 µm.

**Table S1.**
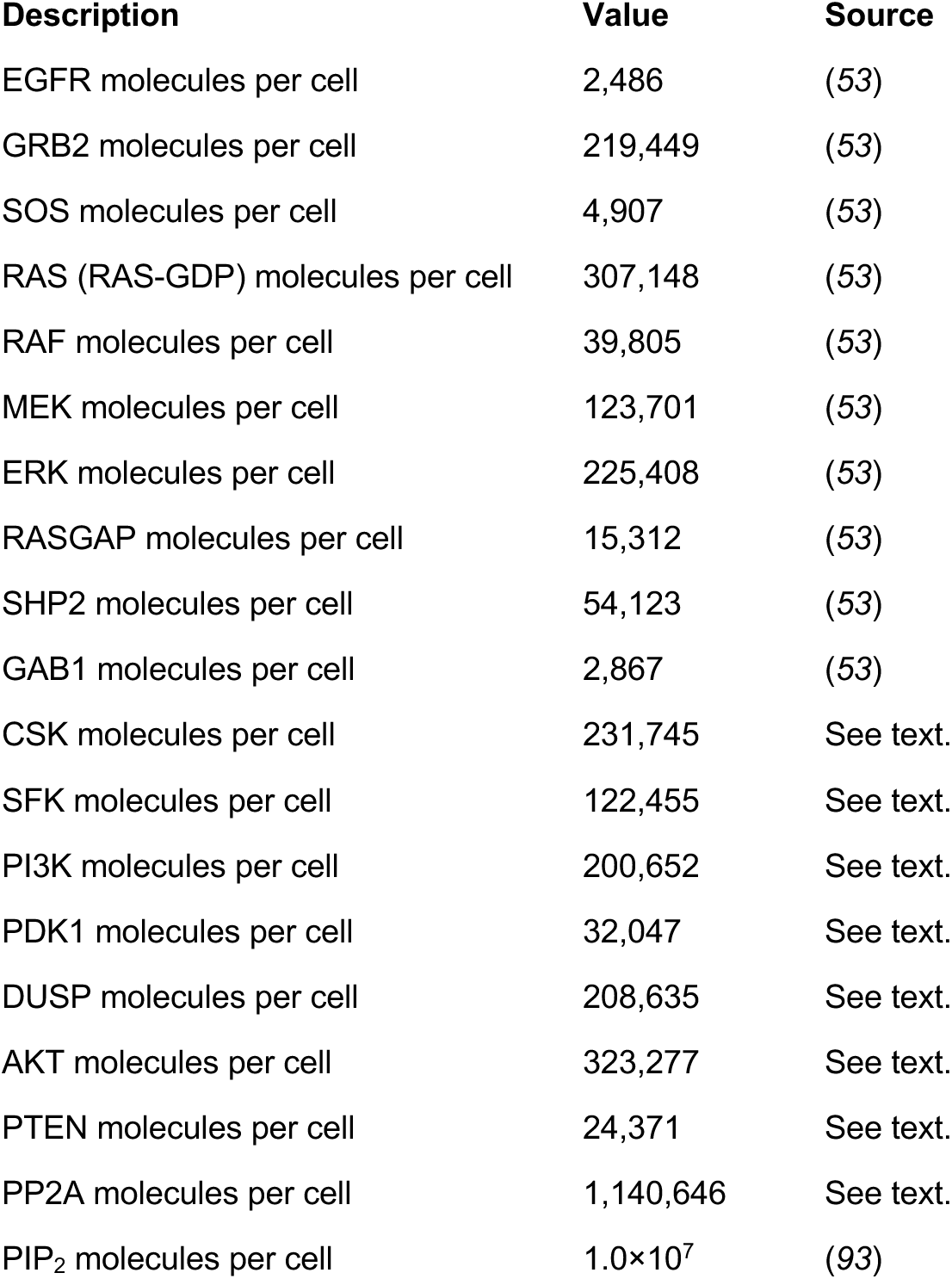
Mechanistic model initial species abundances.

| Description | Value | Source |
| --- | --- | --- |
| EGFR molecules per cell | 2,486 | (53) |
| GRB2 molecules per cell | 219,449 | (53) |
| SOS molecules per cell | 4,907 | (53) |
| RAS (RAS-GDP) molecules per cell | 307,148 | (53) |
| RAF molecules per cell | 39,805 | (53) |
| MEK molecules per cell | 123,701 | (53) |
| ERK molecules per cell | 225,408 | (53) |
| RASGAP molecules per cell | 15,312 | (53) |
| SHP2 molecules per cell | 54,123 | (53) |
| GAB1 molecules per cell | 2,867 | (53) |
| CSK molecules per cell | 231,745 | See text. |
| SFK molecules per cell | 122,455 | See text. |
| PI3K molecules per cell | 200,652 | See text. |
| PDK1 molecules per cell | 32,047 | See text. |
| DUSP molecules per cell | 208,635 | See text. |
| AKT molecules per cell | 323,277 | See text. |
| PTEN molecules per cell | 24,371 | See text. |
| PP2A molecules per cell | 1,140,646 | See text. |
| PIP<sub>2</sub> molecules per cell | 1.0×10<sup>7</sup> | (93) |

**Table S2.** Nominal and initial parameter values.

| **Description** | **Value** | **Units** | **Source** |
| --- | --- | --- | --- |
| EGF binding to EGFR, forward (plasma membrane) | $7.62 \times 10^{-1}$ | $\mu\text{m}^3 \text{ molec}^{-1} \text{ min}^{-1}$ | Calculated |
| EGF binding to EGFR, forward (endosomes) | $4.14 \times 10^{-1}$ | $\mu\text{m}^3 \text{ molec}^{-1} \text{ min}^{-1}$ | Calculated |
| EGF-EGFR binding affinity | 1.6 | nM | (114) |
| EREG binding to EGFR, forward (plasma membrane) | $7.66 \times 10^{-1}$ | $\text{molec}^{-1} \text{ min}^{-1} \text{ cell}$ | Calculated |
| EGF binding to EGFR, forward (endosomes) | $4.16 \times 10^{-1}$ | $\text{molec}^{-1} \text{ min}^{-1} \text{ cell}$ | Calculated |
| EREG-EGFR binding affinity | 76 | nM | Calculated |
| EGFR dimerization, forward (plasma membrane) | $3.63 \times 10^{-3}$ | $\text{molec}^{-1} \text{ min}^{-1} \text{ cell}$ | Calculated |
| EGFR dimerization, forward (endosomes) | $1.95 \times 10^{-3}$ | $\text{molec}^{-1} \text{ min}^{-1} \text{ endosome}$ | Calculated |
| EGF-bound EGFR dimer affinity | $4.8 \times 10^2$ | $\text{molec cell}^{-1}$ | (5, 119) |
| EREG-bound EGFR dimer affinity | $2.88 \times 10^4$ | $\text{molec cell}^{-1}$ | (5, 119) |
| EGFR phosphorylation | $1.3 \times 10^1$ | $\text{min}^{-1}$ | (99) |
| EGFR dephosphorylation (plasma membrane) | $1.7 \times 10^0$ | $\text{min}^{-1}$ | (13) |
| EGFR dephosphorylation (endosomes) | $1.5 \times 10^0$ | $\text{min}^{-1}$ | (13) |
| SFK activation | $1.3 \times 10^1$ | $\text{molec}^{-1} \text{ min}^{-1} \text{ cell}$ | (99) |
| SFK inactivation, CSK-catalyzed | $1.0 \times 10^1$ | $\text{molec}^{-1} \text{ min}^{-1} \text{ cell}$ | (120, 121) |
| GRB2 binding to EGFR, forward | $1.6 \times 10^0$ | $\text{molec}^{-1} \text{ min}^{-1} \text{ cell}$ | (122) |
| GRB2 binding to EGFR, reverse | $4.6 \times 10^2$ | $\text{min}^{-1}$ | (122) |
| GAB1 binding to GRB2, forward | $1.0 \times 10^{-1}$ | $\text{molec}^{-1} \text{ min}^{-1} \text{ cell}$ | (27) |
| GAB1 binding to GRB2, reverse | $6.0 \times 10^1$ | $\text{min}^{-1}$ | (27) |
| PIP<sub>3</sub>-bound GAB1 binding to EGFR:GRB2 | $1.0 \times 10^{-1}$ | $\text{min}^{-1}$ | (27) |
| PIP<sub>3</sub>-bound GRB2 binding to EGFR | $1.0 \times 10^{-1}$ | $\text{min}^{-1}$ | (27) |
| GAB1 phosphorylation, SFK-catalyzed | $1.0 \times 10^{-4}$ | $\text{molec}^{-1} \text{ min}^{-1} \text{ cell}$ | (29) |
| GAB1 phosphorylation, EGFR-catalyzed | $6.0 \times 10^0$ | $\text{min}^{-1}$ | (20) |
| GAB1 dephosphorylation | $9.5 \times 10^0$ | $\text{min}^{-1}$ | (29) |
| SHP2 binding to phosphorylated GAB1, forward | $1.6 \times 10^0$ | $\text{molec}^{-1} \text{ min}^{-1} \text{ cell}$ | (122, 123) |
| SHP2 binding to phosphorylated GAB1, reverse | $4.6 \times 10^2$ | $\text{min}^{-1}$ | (122, 123) |
| SOS binding to GRB2, forward | $6.0 \times 10^{-4}$ | $\text{molec}^{-1} \text{ min}^{-1} \text{ cell}$ | (20) |
| SOS binding to GRB2, reverse | $6.0 \times 10^0$ | $\text{min}^{-1}$ | (20) |
| PI3K binding to phosphorylated GAB1, forward | $6.0 \times 10^{-4}$ | $\text{molec}^{-1} \text{ min}^{-1} \text{ cell}$ | (20) |
| PI3K binding to phosphorylated GAB1, reverse | $6.0 \times 10^0$ | $\text{min}^{-1}$ | (20) |
| PIP<sub>3</sub> phosphorylation | $6.0 \times 10^{-3}$ | $\text{min}^{-1}$ | (20) |
| PDK1 binding to PIP<sub>3</sub>, forward | $6.0 \times 10^{-4}$ | $\text{molec}^{-1} \text{ min}^{-1} \text{ cell}$ | (20) |
| PDK1 binding to PIP<sub>3</sub>, reverse | $6.0 \times 10^0$ | $\text{min}^{-1}$ | (20) |
| AKT binding to PIP<sub>3</sub>, forward | $6.0 \times 10^{-4}$ | molec<sup>-1</sup> min<sup>-1</sup> cell | (20) |
| AKT binding to PIP<sub>3</sub>, reverse | $6.0 \times 10^0$ | min<sup>-1</sup> | (20) |
| AKT phosphorylation | $6.0 \times 10^1$ | min<sup>-1</sup> | (20) |
| AKT dephosphorylation, PP2A-catalyzed | $6.0 \times 10^{-4}$ | molec<sup>-1</sup> min<sup>-1</sup> cell | (30) |
| PTEN binding to PIP<sub>3</sub>, forward | $6.0 \times 10^{-4}$ | molec<sup>-1</sup> min<sup>-1</sup> cell | (20) |
| PTEN binding to PIP<sub>3</sub>, reverse | $6.0 \times 10^0$ | min<sup>-1</sup> | (20) |
| PIP<sub>3</sub> dephosphorylation, PTEN-catalyzed | $6.0 \times 10^{-1}$ | min<sup>-1</sup> | (20) |
| RAS GDP-GTP exchange, SOS-catalyzed | $6.0 \times 10^{-5}$ | molec<sup>-1</sup> min<sup>-1</sup> cell | (30) |
| RASGAP binding to RAS-GTP, forward | $6.0 \times 10^{-4}$ | molec<sup>-1</sup> min<sup>-1</sup> cell | (20) |
| RASGAP binding to RAS-GTP, reverse | $6.0 \times 10^0$ | min<sup>-1</sup> | (20) |
| Intrinsic RAS-catalyzed GTP hydrolysis | $4.02 \times 10^2$ | min<sup>-1</sup> | (107) |
| RASGAP-catalyzed RAS-GTP hydrolysis | $2.82 \times 10^{-2}$ | min<sup>-1</sup> | (107) |
| SHP2-catalyzed RASGAP-RAS disassociation | $6.0 \times 10^{-5}$ | molec<sup>-1</sup> min<sup>-1</sup> cell | (30) |
| RAF phosphorylation, RAS-GTP-catalyzed | $6.0 \times 10^{-5}$ | molec<sup>-1</sup> min<sup>-1</sup> cell | (30) |
| RAF dephosphorylation, PP2A-catalyzed | $6.0 \times 10^{-4}$ | molec<sup>-1</sup> min<sup>-1</sup> cell | (30) |
| MEK phosphorylation, pRAF-catalyzed | $6.0 \times 10^{-5}$ | molec<sup>-1</sup> min<sup>-1</sup> cell | (30) |
| MEK dephosphorylation, PP2A-catalyzed | $6.0 \times 10^{-4}$ | molec<sup>-1</sup> min<sup>-1</sup> cell | (30) |
| ERK phosphorylation, pMEK-catalyzed | $6.0 \times 10^{-5}$ | molec<sup>-1</sup> min<sup>-1</sup> cell | (30) |
| ERK dephosphorylation, DUSP-catalyzed | $6.0 \times 10^{-4}$ | molec<sup>-1</sup> min<sup>-1</sup> cell | (30) |
| SOS negative feedback phosphorylation, pERK-catalyzed | $6.0 \times 10^{-5}$ | molec<sup>-1</sup> min<sup>-1</sup> cell | (30) |
| SOS negative feedback dephosphorylation | $6.0 \times 10^{-2}$ | min<sup>-1</sup> | (30) |
| RAF negative feedback phosphorylation, pERK-catalyzed | $6.0 \times 10^{-5}$ | molec<sup>-1</sup> min<sup>-1</sup> cell | (30) |
| RAF negative feedback site dephosphorylation, PP2A-catalyzed | $6.0 \times 10^{-5}$ | molec<sup>-1</sup> min<sup>-1</sup> cell | (30) |
| MEK negative feedback phosphorylation, pERK-catalyzed | $6.0 \times 10^{-5}$ | molec<sup>-1</sup> min<sup>-1</sup> cell | (30) |
| MEK negative feedback site dephosphorylation, PP2A-catalyzed | $6.0 \times 10^{-5}$ | molec<sup>-1</sup> min<sup>-1</sup> cell | (30) |
| GRB2-bound EGFR endocytosis | $2.0 \times 10^{-1}$ | min<sup>-1</sup> | (108) |
| Ligand-bound EGFR endosomal exit | $4.0 \times 10^{-2}$ | min<sup>-1</sup> | (108) |
| Ligand-bound EGFR recycling fraction | $5.0 \times 10^{-1}$ | N/A | (108) |
| Ligand-unbound EGFR endosomal exit | $4.0 \times 10^{-2}$ | min<sup>-1</sup> | (108) |
| Ligand-unbound EGFR recycling fraction | $8.0 \times 10^{-1}$ | N/A | (108) |

## ACKNOWLEDGEMENTS

We thank Dr. Dan Freed (Chordoma Foundation), Dr. Anatoly Kiyatkin (Yale School of Medicine), and Dr. Mark Lemmon (Yale School of Medicine) for helpful technical discussions. Figure 1 was created with BioRender.com.

## Author contributions

Conceptualization, P.J.M and M.J.L; Methodology, P.J.M., S.L., and M.J.L; Software, P.J.M.; Formal Analysis, P.J.M. and S.L.; Investigation, P.J.M. and S.L.; Visualization, P.J.M. and S.L.; Writing – Original Draft, P.J.M., S.L., and M.J.L.; Funding Acquisition, P.J.M. and M.J.L.

